# STING promotes homeostatic maintenance of tissues and confers longevity with aging

**DOI:** 10.1101/2024.04.04.588107

**Authors:** Jacob W. Hopkins, Katherine B. Sulka, Machlan Sawden, Kimberly A. Carroll, Ronald D. Brown, Stephen C. Bunnell, Alexander Poltorak, Albert Tai, Eric R. Reed, Shruti Sharma

## Abstract

Local immune processes within aging tissues are a significant driver of aging associated dysfunction, but tissue-autonomous pathways and cell types that modulate these responses remain poorly characterized. The cytosolic DNA sensing pathway, acting through cyclic GMP-AMP synthase (cGAS) and Stimulator of Interferon Genes (STING), is broadly expressed in tissues, and is poised to regulate local type I interferon (IFN-I)-dependent and independent inflammatory processes within tissues. Recent studies suggest that the cGAS/STING pathway may drive pathology in various *in vitro* and *in vivo* models of accelerated aging. To date, however, the role of the cGAS/STING pathway in physiological aging processes, in the absence of genetic drivers, has remained unexplored. This remains a relevant gap, as STING is ubiquitously expressed, implicated in multitudinous disorders, and loss of function polymorphisms of STING are highly prevalent in the human population (>50%). Here we reveal that, during physiological aging, STING-deficiency leads to a significant shortening of murine lifespan, increased pro-inflammatory serum cytokines and tissue infiltrates, as well as salient changes in histological composition and organization. We note that aging hearts, livers, and kidneys express distinct subsets of inflammatory, interferon-stimulated gene (ISG), and senescence genes, collectively comprising an immune *fingerprint* for each tissue. These distinctive patterns are largely imprinted by tissue-specific stromal and myeloid cells. Using cellular interaction network analyses, immunofluorescence, and histopathology data, we show that these immune fingerprints shape the tissue architecture and the landscape of cell-cell interactions in aging tissues. These age-associated immune fingerprints are grossly dysregulated with STING-deficiency, with key genes that define aging STING- sufficient tissues greatly diminished in the absence of STING. Changes in immune signatures are concomitant with a restructuring of the stromal and myeloid fractions, whereby cell:cell interactions are grossly altered and resulting in disorganization of tissue architecture in STING-deficient organs. This altered homeostasis in aging STING-deficient tissues is associated with a cross-tissue loss of homeostatic tissue-resident macrophage (TRM) populations in these tissues. *Ex vivo* analyses reveal that basal STING-signaling limits the susceptibility of TRMs to death-inducing stimuli and determines their *in situ* localization in tissue niches, thereby promoting tissue homeostasis. Collectively, these data upend the paradigm that cGAS/STING signaling is primarily pathological in aging and instead indicate that basal STING signaling sustains tissue function and supports organismal longevity. Critically, our study urges caution in the indiscriminate targeting of these pathways, which may result in unpredictable and pathological consequences for health during aging.

**HIGHLIGHTS:**

a. Aging tissues are associated with tissue-autonomous *immune* fingerprints, primarily driven by interactions of tissue stromal and myeloid populations.
b. STING shapes these immune fingerprints of aging tissues in unexpected ways.
c. Loss of STING alters the location, numbers, and viability of tissue resident macrophages.
d. STING signaling is critical for longer lifespans and maintenance of tissue architecture.

## INTRODUCTION

During aging, tissue-associated changes do not proceed in step with shifts in systemic markers of inflammation ^1^. Tissue-autonomous immunity forms the largest fraction of organ-based immune responses; yet studies on aging have overlooked the contribution of cell-autonomous and tissue-intrinsic immunity to

age-associated dysfunction. Recent studies have revealed important roles for the ubiquitously expressed, innate immune DNA-sensor cyclic GMP-AMP synthase (cGAS) and its downstream ER-resident signaling adaptor Stimulator of Interferon Genes (STING) in driving inflammation, type I interferon (IFN-I) responses and cellular senescence in stromal cells like fibroblasts ^2,3^. Based on these roles, it is proposed that cGAS/STING signaling is the mechanistic basis for aging associated dysfunction. This hypothesis has borne out in various transgenic mouse models tied to the aberrant accumulation of cytosolic DNA and its sensing through cGAS/STING driving chronic inflammatory conditions reminiscent of pathological aging^4–6^. However, the mechanistic contribution of the cGAS/STING pathway to lifespans and health spans during physiological aging in multiple tissues and cell types, unmodified by the genetic drivers, remains undetermined. Furthermore, the categorical impact of cGAS/STING-signaling on lifespans in physiologic settings is also unknown. These missing links are critical gaps as a) aging itself is not a disease; b) they fail to fully anticipate or shape the impact of STING agonist/antagonist therapies currently being proposed for use in aging by studies that consider it a pathological condition; and finally, c) at least four well-known loss-of-function (LOF) polymorphic alleles of STING are represented at a frequency of >50% in the human population, underscoring the need to understand how lack of STING-signaling shapes physiological aging processes^7–9^. Here we evaluate the impact of STING-deficiency during physiologic aging on three separate tissues: the heart, liver, and kidneys, critical determinants of healthy aging, as well as on overall longevity and health. We reveal how aging imprints these key tissues with unique inflammatory, interferon-stimulated gene (ISG), and senescence gene signatures that are akin to immune ‘*fingerprints’*. This immunological fingerprinting of tissues is driven primarily by stromal and myeloid compartments, shaping the tissue architecture and landscape of cell:cell interactions as organs age. We uncover that STING signaling contributes in unexpected ways to the generation of these fingerprints, by regulating the expression of these key genes and suppressing the expression of other pro-inflammatory gene-sets. These changes are concomitant with reorganization of the tissue-architecture and changes in cell:cell interactions in the absence of STING. Collectively, these differences result in significantly *shorter* organismal lifespans and perturbed homeostasis systemically as well as in the organs of STING-deficient mice. Mechanistically, STING-deficiency is associated with a significant loss in tissue-resident macrophage populations that have been homeostatically programmed within these specific tissues ^10,11^. STING-signaling stipulates the sensitivity of TRMs to various cell-death stimuli and thus also regulates their survival, cell:cell interaction and spatial localization with aging. cGAS/STING signaling are considered the mechanistic drivers of pathological aging. Here we reveal data that fundamentally challenges this concept and instead reveals that cGAS/STING serve as a critical homeostatic checkpoint during aging, and a vital longevity determinant. Our study thus emphasizes a need to carefully evaluate how STING therapeutics are utilized and especially how they interface with any existing loss of function in this pathway.

## RESULTS

### TISSUE AUTONOMOUS INFLAMMATORY SIGNATURES DRIVEN BY LOCAL STROMAL AND MYELOID CELLS IMPRINT AGING TISSUES

Recirculating innate immune markers and tissue inflammation remain weakly linked. Thus, to explicate this relationship, we measured serum cytokines and chemokines at 3 months old (mo), 12mo, and 24mo. Notable among these signatures were the elevated profiles of key innate inflammatory cytokines, but also broadly immunoregulatory cytokines and chemokines. (**Figure 1A**). Concomitant with these increased serum-cytokine signatures, histological parameters of tissue disorganization and dysfunction were also apparent in aged tissues (**Figure S1A**). We primarily evaluated changes in hearts, livers, and kidneys, as previous aging studies have highlighted the importance of these tissues in shaping the trajectory of healthy or unhealthy aging ^12^. In aged hearts, livers, and kidneys, we note a general disruption of the tissue architecture associated with increased immune infiltrates surrounding the larger vasculature, with specific instances of stromal reorganization and tubular inflammation in the hearts and kidneys respectively (**Figure S1A**). Various theories of aging have suggested that peripheral immune changes, such as elevated cytokine signatures in plasma/sera, directly impact the aging of tissues themselves and are reliable markers of tissue dysfunction ^12^. Although recent studies note inflammatory changes in tissue gene expression profiles, the relationship between systemic and tissue-associated markers of inflammation remains unexplored. To investigate this gap, we leveraged unbiased single cell-RNA sequencing (scRNAseq) of the three distinct organs (heart, liver, and kidney) over 4 time points (**Figure S1B**). The three organs can reliably be segregated by cell type, tissue type and age (**Figure 1B and S1C-D**).

**Figure 1:**
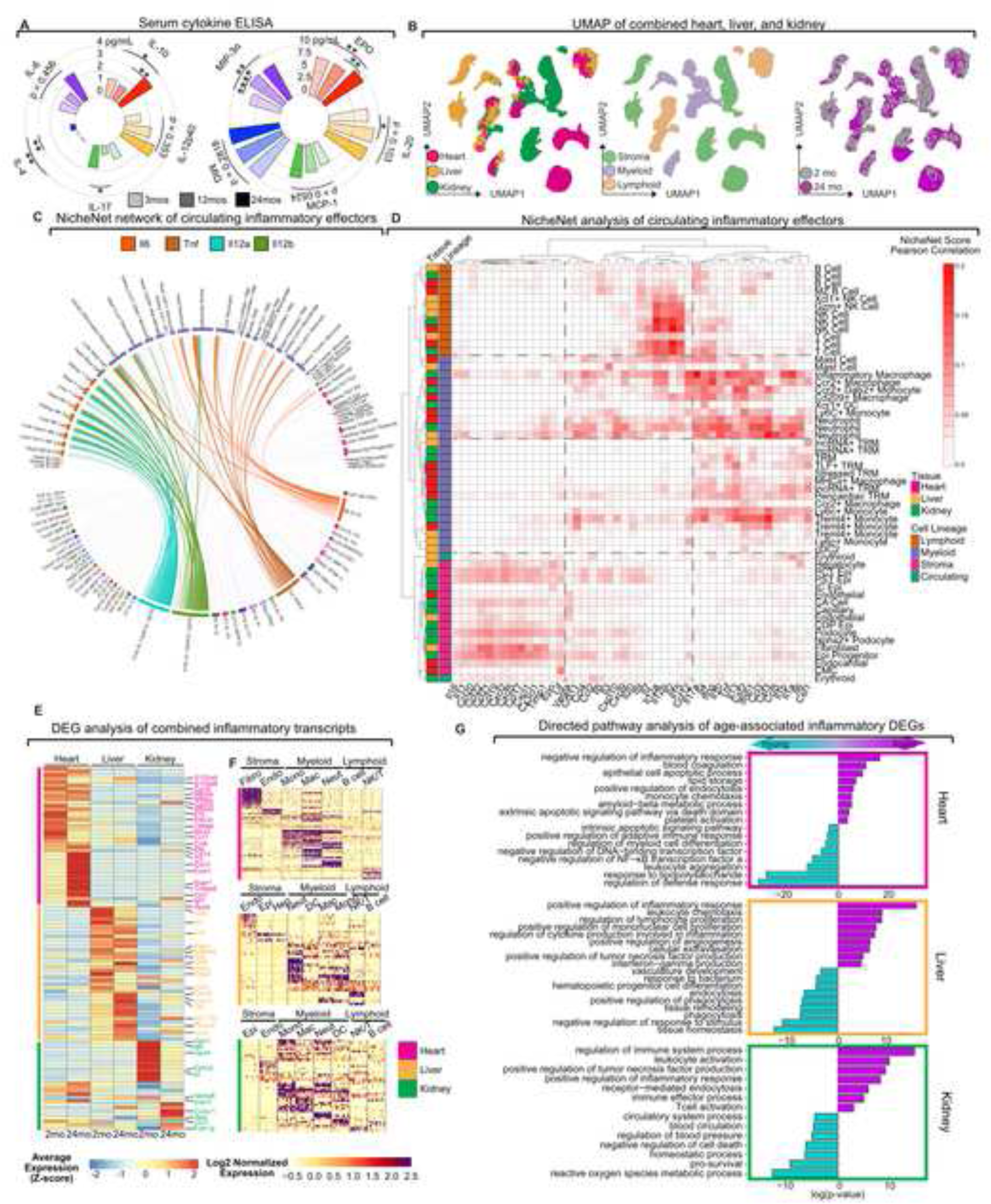
Aging stromal and myeloid cells shape tissue intrinsic inflammatory fingerprints. **(A)** Radial bar chart depicting serum concentrations (pg/mL) of inflammatory cytokines and chemokines (immunokines) from 3, 12, and 24-mo WT mice measured by multiplex ELISA (n = 4-5 mice). **(B)** Combined Uniform Manifold Approximation and Projection (UMAP) of tissue-associated cells from 2- and 24-mo heart, liver, and kidneys. Clusters are highlighted by tissue, lineage, and age (left to right, annotated as shown). **(C, D)** NicheNet analysis indicating the ligand– target regulatory potential elicited in 24-mo WT mice cell types by the immunokines assayed in (a). **(c)** Circos plot visualization of active ligand–target links between cells and immunokines assayed in (a). *Il6*, *Tnf*, *Il12a*/*b* highlighted. **(D)** Heatmap representation of mean NicheNet scores (based on Pearson correlation coefficients) of ligand–target regulatory potential scores across tissue-associated cells from the heart, liver, and kidney. Tissue and cell lineage are annotated. **(E)** Differential gene expression (DGE) analysis of top 25 inflammatory marker genes in the combined datasets of all three tissues and both ages (2 and 24mo). Bonferroni adjusted p-value < 0.1, for 2mo and 24mo WT samples from each tissue (hot pink, heart; gold, liver; green, kidney). Inflammatory genes compiled from gene ontology (GO):0006954. Gene expression scaled for row maximum. **(F)** Single cell expression levels of top inflammatory marker genes, Bonferroni adjusted p-value < 0.1, by cell type and tissue of origin from the combined dataset. Fibro, fibroblast; Endo, Endothelial; Epi, epithelial; Hep, hepatocyte; Mono, monocyte; Mac, macrophage; Neut, neutrophil; DC, dendritic cell; B, B cell; NK/T, NKT cell/T cell). **(G)** Bar chart depicting top GO pathways enriched in inflammatory DEGs during youth and aging. Pathways positively correlated with age in magenta, pathways negatively correlated with age (positive with youth) in cyan (hot pink, heart; gold, liver; green, kidney). Statistics calculated using ANOVA (a), or Wilcoxon Rank Sum test (e, f). \**p ≤ 0.05*; \*\**p ≤ 0.01*; \*\*\**p ≤ 0.001*.

To evaluate the extent to which the observed tissue pathology in distinct cellular components may be driven by circulating chemokines or cytokines, we sought to overlay our serum cytokine profile upon the scRNAseq dataset by leveraging the NicheNet analytical tool ^13^. The NicheNet platform models ligand– receptor interactions in cells by combining their scRNAseq expression data with existing knowledge of signaling and gene regulatory networks. In this instance, we impute serum cytokines as our ligands and assign the overlap between gene-expression data in cells from the three aging tissues and target gene-modules downstream of these cytokines as an enrichment score. We note an enrichment of genes that map to well-described, cytokine signaling modules, confirming that the interactions between cytokines and their cognate receptors are observable in all three tissue datasets (**Figure 1C-D**). Indeed, while signaling components downstream of key serum cytokines are established in all three tissues, the identified cells responding to these key cytokines are often the same in these distinct tissues **(Figure 1C)**. For instance, IL- 12p40 or p70 gene programs are identified in NK/T cells, while IL-6/TNFα appear to mostly impact transcriptional modules of innate immune myeloid cells in the three tissues (**Figure 1C**). These data suggest that the expression profile of receptors on specific cell types in each tissue may limit or determine the impact of circulating cytokines. Importantly, a broader view of the relative expression of these cytokine signatures suggests that peripheral cytokine signaling leads to comparable downstream signaling consequences for similar cell types in a tissue independent manner **(Figure 1D)**. This is overwhelmingly true of immune cells and holds to broad groups of stromal/parenchymal populations like epithelial cells, endothelial cells, and fibroblasts, which respond in analogous ways to the cyto- or chemokines by upregulating highly similar gene expression modules (**Figure 1D**). These data suggest that tissue immune signatures were likely to be highly similar in different tissues as they would be the product of similar cytokine signaling on cell types found in all three tissues.

To then test if aging tissues present with similar gene expression profiles of inflammatory genes (INF), we first analyzed the differentially expressed genes (DEGs) in each tissue with age. Similar to previous reports, we can successfully separate cellular identities in tissues using the expression of core genes (Figure **S2A-C**) ^14,15^. We also note that the enrichment of DEGs in aged tissues is equally distributed between stromal, myeloid, and lymphoid compartments (**Figure S2B-C, D**). A comprehensive, comparative understanding of immune signatures and how they segregate by age and tissue type is fundamental to further enquiry into the relationship between tissue immune signatures and recirculating serum cytokines/chemokines. Thus, we next queried the expression of 370 immune genes (drawn from GO-annotation pathways and hallmark datasets) in the three organs with age (**Figure 1E and Supplemental Information 1**) ^16^. Surprisingly, we note a distinctive immune signature for each tissue, which we refer to as an ‘immune fingerprint’ (**Figure 1E**). These immune fingerprints were altered in each organ with age but remained distinct between organs (**Figure 1E**). Whilst there were instances of common genes expressed in all three tissues (*Ccr2, Vamp8, Ccr7, Apoe*, etc.), these genes did not comprise the majority of the age-related changes in gene expression. Deepening our understanding of this unique profile – we find that these immune fingerprints are enriched primarily in stromal/parenchymal compartments and the tissue associated myeloid subsets in all three tissues, but relatively little of the signature is derived from the lymphoid component (**Figure 1F**). Pathway analysis of inflammatory genes enriched in young and aged tissues, respectively, confirms that there is a relative enrichment of classically inflammatory gene sets in the aged tissues, whereas young tissues instead are enriched in immune genes with a bias towards regulation of inflammatory and tissue homeostasis **(Figure 1G)**. Collectively, these data suggest that tissue immune profiles are highly unique to each organ and cannot solely be attributed to the more invariant effect of immune signaling from circulating cytokines. Furthermore, the disproportionate contribution of the stromal and myeloid compartments to these signatures underscores the importance of cell-autonomous immune responses in shaping local tissue immune fingerprints.

### INTERFERON SIGNALING AND SENESCENCE ARE CORE ATTRIBUTES OF SUPPORTIVE AND TISSUE-IMMUNE ELEMENTS IN AGING TISSUES

In order to understand the composition of the immune profile in aging tissues, we performed pathway analysis on genes selectively upregulated in aged cell types of each tissue relative to young (**Figure 2A and Supplemental Information 2**). Notably, this analysis emphasizes the enrichment of immune activation and chemotactic/migratory signatures in aging cell types, confirming data presented in **Figure 1G**. Aging also enacts/promotes significant enrichment of stress-related pathways, cell-death signaling, cytoskeletal remodeling and critically, two core inflammatory gene sets: interferon stimulated genes (ISGs) and senescence genes (SEN) (**Figure 2A** and **Supplemental Information 1**). ISGs in our pathway analysis match those previously identified as IFN-I inducible genes (**Figure S3A**) ^17^. Critically, though highly relevant in chronic diseases, the role of IFN-I in tissue immunity during physiological aging is largely unexplored. IFN-I signaling is thought to result in the upregulation of ≥700 ISGs in a temporal and cell-specific manner ^17^. Gene expression analysis identified 575 ISGs in our scRNAseq dataset and reveals their overall upregulation with age. Importantly, like immune genes, distinct clusters of ISGs define each tissue, and are further modified with age, identifying an ‘ISG fingerprint’ for resting and aging tissues (**Figure 2B**). For instance, in the aging heart *Socs3*, *Ifitm3*, *Ccl4* are among the key ISGs differentially expressed compared to the younger hearts (*Isg15, Mxd1, Vps37b)* (**Figure 2B**). These ISGs are different from those highly expressed in the aging livers (*Gzmb, Irf8, Id2*) and kidneys (*Bst2, Ifi44, C1qc*) (**Figure 2B**). As with inflammatory signatures, ISGs were primarily enriched within stromal-myeloid cell types, although the composite ISG signature is, in part, also derived from lymphoid cells, particularly in the liver (**Figure 2C).**

**Figure 2:**
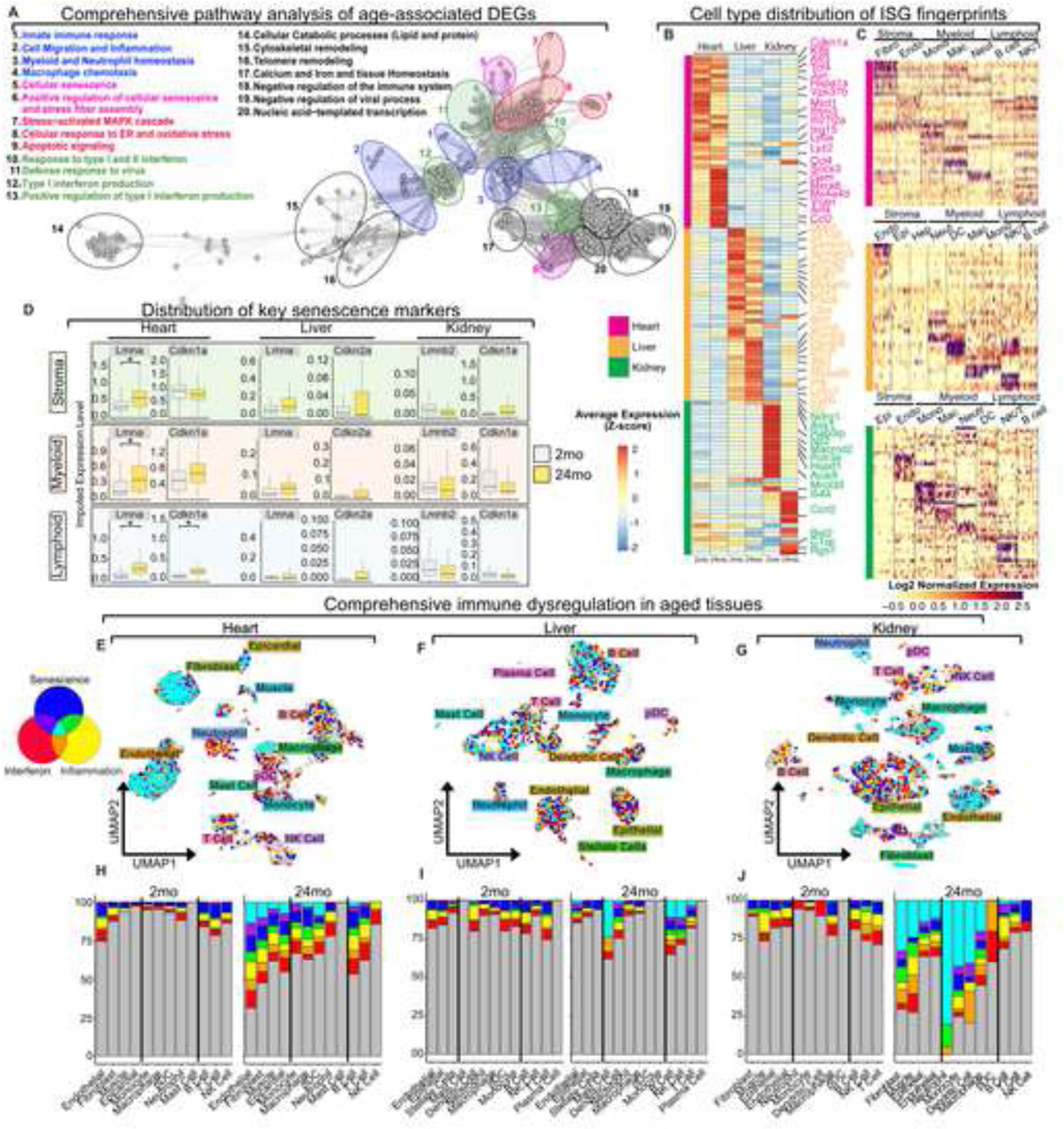
Immune fingerprints in aging tissues are enriched for interferon and senescence signatures. (A) GO pathway network analysis for all DEGs upregulated with age per cell type. GO terms were summarized by semantic similarity with Revigo. Nodes represent individual pathways upregulated in aged cell types, connecting edges represent relative semantic similarity. Blue, innate immune response; pink, senescence; green, interferon responsiveness; red, cellular stress; black, cellular function**. (B)** Expression of top 25 interferon stimulated marker genes (Bonferroni adjusted p-value < 0.1) for 2mo and 24mo WT tissues (heart, hot pink; liver, gold; kidney, green). **(C)** Single cell relative expression levels of top interferon stimulated marker genes per cell type separated by tissue of origin. (Fibro, fibroblast; Endo, endothelial; Epi, epithelial; Hep, hepatocyte; Mono, monocyte; Mac, macrophage; Neut, neutrophil; DC, dendritic cell; B, B cell; NK/T, NKT cell/T cell) **(D)** MAGIC imputed expression levels of key senescence associated genes by cell lineage and tissue. Box plots depict median expression level, first, third quartiles, and range. (Boxplots: grey, 2mo; yellow, 24mo). **(E-G)** UMAP with maximal relative GSVA enrichment per cell of inflammatory (yellow), interferon response (red), or senescence associated (blue) metagenes in the heart **(E)** liver **(F)** and kidney **(G)**. Cells relatively enriched in all three immune pathways (polypositive, PP) in cyan. **(H-J)** Stacked bar plots depict relative distribution of cells enriched for unique or overlapping immune metagenes, including proportion of PP cells, in 2mo and 24mo samples. Heart **(H)**, liver **(I)**, kidney **(J)**. Statistics calculated using a Wilcoxon Rank Sum test. \**p ≤ 0.05*; \*\**p ≤ 0.01*; \*\*\**p ≤ 0.001*.

Senescence is considered a hallmark of aging tissues and may contribute to tissue immune profiles via multiple inflammatory genes and ISGs that make up the Senescence-Associated Secretory Phenotype (SASP)^18,19^. Previous studies have relied primarily on *ex vivo* analyses or reporters of key cell cycle regulators *Cdkn1a* (p21) and *Cdkn2a* (p16), to illuminate the relationship between aging and senescence, but have yielded conflicting results^20^. While recent years have seen *ad hoc* use of scRNAseq on select tissue cells to investigate senescence, a comprehensive single cell resolution map of senescence genes in physiologically aged tissues is lacking^21^. With our large, multi-tissue scRNAseq dataset, we sought to evaluate key senescence markers to understand their fluctuation with age. Our analysis reveals that the expression of senescence markers such as *Cdkn2a/1a* (p16/p21) or *Lmnb1* is neither consistent nor temporally coordinated in the three different aging tissues (**Figure 2D and S3B-C**). Furthermore, different elements of the tissue, such as stromal, myeloid, and lymphoid compartments, contribute in different ways to the development of senescence profiles in tissues (**Figure 2D**). Distinct senescence genes reflecting the SASP phase are notably altered in the stromal compartments of the heart (*Lmna, Timp2, Cd47, Ctgf, Igfbp4, and Cxcl1*), liver (*Cd47, Ctgf*), and kidneys (*Il1a, Cd47, Ccl5*) (**Figure 2D, S3C**). We also note similar differences in the myeloid compartment, with all tissue-associated myeloid cells upregulating a large contingent of canonical senescence and SASP genes. Myeloid cells in the aging heart preferentially upregulate *Il1b, Timp2, Lmna*, *Igfbp4*, *Ccl2*; whilst aged liver myeloid cells upregulate other markers of SASP including *Igfbp4* and kidney myeloid cells express yet other SASP genes (*Mmp9, Ccl4,* Ccl3) (**Figure S3D**). Finally, the lymphoid compartment also contributes to senescent profiles in the heart (*Cdkn1a, Timp2, Cd47, Lmna, Ccl5*), liver (*Ccl4, Igfb4, Ccl5, Ccl3*) and kidney (*Ccl4, Ccl5)* (**Figure 2D and S3C**). Altogether, these data reveal that the immune fingerprints of young and especially aged tissues are a complex amalgamation of many innate immune gene-sets, or metagenes, that collectively shape tissue behavior during aging.

Whether distinct cellular components generated the cumulative tissue immune profiles noted above, was tested using Gene Set Variation Analysis (GSVA) to reduce the dimensionality of these extensive signatures into a singular scoring system. Here, calculating individual “activity scores” for each geneset allows single cell quantification of the **relative** enrichment of its genes, accounting for the breadth and depth of expression of individual genes within the signature **(Figure 2E-G, S4A, Supplemental Information 1)**. This enables a new, clearer understanding of the relative distribution of these immune signatures with age, as we can reliably measure the activity score of each signature rather than relative proportions of tens or hundreds of cells. Using the mean activity score of clusters from the 2mo dataset as a reference, we were able to normalize the activity scores of different cell types for each signature to best understand the distribution and directionality of these signatures with age. These data reveal that while the individual signatures are stochastically distributed in the tissues with age, the overlap of the three metagene signatures (INF, ISG and SEN; referred to here as polypositive; PP) is only apparent in aged tissues and further in highly specific stromal and myeloid cell populations (**Figure 2E-J and S4B-D**). Whilst overall cellularity may be decreased with age (**Figure S4B-D**), the proportion of PP cells increases with age, as do cells enriched for more than one signature (double positive; DP) (**Figure 2H-J**). Although the cellular compartments that dominantly contribute to the PP signature are primarily stromal-myeloid in nature, the genes upregulated in similar cell groups in different tissues is unique. For instance, *Ets1, Ptgs2, and Cxcr5* are specifically upregulated in cardiac fibroblasts, whereas renal fibroblasts preferentially express *Ifi2712a and Ccl12* (**Figure S4E-G**). These data are the first to highlight how only a few cell types contribute overwhelmingly to an organ’s immune fingerprint with aging. Furthermore, as only specific cell types drive the *coordinated* expression of several immune pathways, their critical roles in tissue fingerprinting may be shaped by innate immune pathways selectively expressed within these cells.

### THE STING-PATHWAY RE-SHAPES THE TISSUE IMMUNE LANDSCAPE

The enrichment of INF, ISG and SEN metagenes in the cells of stromal-myeloid lineages, implicates innate immune mechanisms that can shape all three signatures in determining these fingerprints with age. Contrary to other innate pattern-recognition receptors (PRRs), recent evidence suggests cGAS/STING activation may regulate the expression of senescence markers, in addition to shaping inflammation and ISG expression, thus promoting unhealthy aging^3^. First, we assessed how STING signaling contributes to the three “metagene” fingerprints in aging tissues. Initial cell subset analysis of tissues from adult (2mo) and aged (24mo) STING^-/-^ mice reveals that overall cellular distribution and annotation in the tissues is akin to WT mice (**Figure 3A)**. Previous studies reveal how STING drives inflammatory genes like IL-6 and TNFα in addition to key ISGs and SASP genes^22^. Thus, an overall *loss* of the three metagene signatures would be predicted in the absence of STING. Indeed, we note that inflammatory markers prevalent in WT tissues like *Saa3, Cxcl2, Ccl6, C3ar1*; ISGs like *Bst2, Ifitm3, Slamf7, Fn1, Ifi205, Ifit2/3 Mx1*; and SEN genes *Cdkn2a, Calm1/2/3, Atr,* are decreased in aged STING^-/-^ hearts, livers, and kidneys (**Figure 3B**). However, comprehensive comparative analysis of cumulative INF, ISG, and SEN signatures in aging STING^-/-^ tissues reveals that the metagene signatures are not merely lost or lower but are categorically *different* from the age-matched WT (**Figure 3B)**. Indeed, highly proinflammatory genes like *Cd14, Nfkb, Pparg, Cd68, Il1b, Akt, Pycard*; ISGs, *Irf1, Ptgs2, Irf8, Stat1, Plac8, Casp8* and SEN genes*, Cdkn1a, H2-Q proteins, Pten, Tgfbr1/2, Lmna, Akt2, Nfatc1,* are all unexpectedly upregulated in STING^-/-^ tissues.

**Figure 3:**
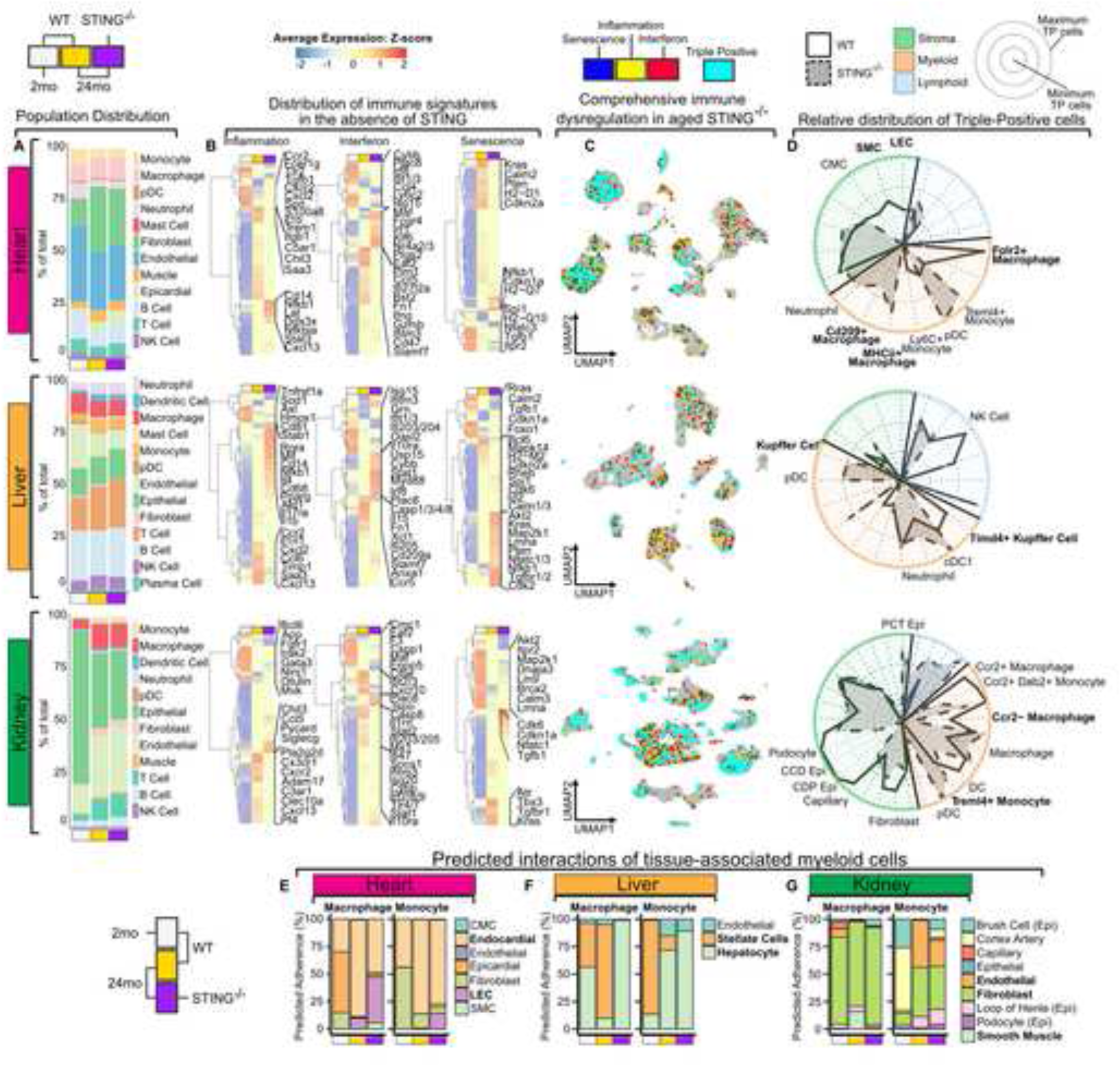
An altered immune landscape in the absence of STING. (A) Stacked bar plots indicate relative distribution of indicated cell types in tissue-specific data sets. Age and genotype are indicated below (2mo WT, grey; 24mo WT, yellow; 24mo STING^-/-^, purple) **(B)** Relative average expression of top inflammatory, interferon, and senescence associated DEGs in 2mo and 24mo WT and 24mo STING^-/-^ tissues identified by Bonferroni adjusted p-value < 0.1. Age and genotype indicated above are consistent with (a). **(C)** UMAP representation of maximal enrichment of all three immune metagenes in 24mo STING^-/-^ tissues **(D).** Radar plot quantification of relative enrichment of polypositive cells (solid lines: WT; dashed lines: STING^-/-^) in cell lineage subclusters. Key populations annotated for clarity (green, stroma; orange, myeloid; blue, lymphoid). **(E)** Stacked bar plots of predicted adherence between macrophages or monocytes and distinct stromal cell anchors forecast by the RNAMagnet algorithm (Velten Lab). Age and genotype indicated below are the consistent with (a). Differentially expressed genes calculated using a Wilcoxon rank sum test (b).

Plotting the co-occurrence of INF, ISG, and SEN signatures reveals that cell types enriched in all three fingerprints i.e., PP cell types, are different in the aged STING^-/-^ tissues compared to WT controls (**Figures 3C** and **2E-G**). Quantitation of relative enrichment of PP cells in WT vs STING^-/-^ tissues reveals how this distribution shifts (**Figure 3D**). In the aged WT heart, the PP signature is primarily enriched in key stromal populations and a subset of tissue macrophages (Folr2^+^) (**Figure 3D**). However, STING-deficiency shifts the brunt of the signature to multiple myeloid fractions, with significant loss of PP signatures notable in the lymphatic endothelial cells, smooth muscle cells, and Folr2^+^ macrophages (**Figure 3C-D**). Similar redistribution of PP stromal and myeloid populations is also evident in the aging STING^-/-^ livers and kidneys, with a consistently broader set of myeloid cells that become PP in the absence of STING (**Figure 3C-D**). Thus, our data underscore that existing knowledge of STING-dependent regulation of immune genes is incomplete and single-cell analysis combined with broad inquiry in fact reveal the breadth of immune signatures that STING regulates *in vivo*.

We hypothesized that changes to INF, ISG, and SEN signatures in STING^-/-^ tissues likely impact cell-cell interactions in tissues. To this end, we leveraged RNA-magnet analysis which predicts the potential physical interactions between designated anchor populations and target cells based on the expression patterns of cell-surface receptors and cognate ligands^23^. This analysis thus allows us to query interaction probabilities, strengths, and signaling nodes, between annotated cell types in the tissues (**Figure 3E-G**). We focused primarily on myeloid/macrophage subsets as targets and stromal components as anchors, as these were the most signature enriched cell subsets (**Figure 3C-D**). Both macrophages and monocytes from WT and STING^-/-^ hearts preferentially engage a variety of endothelial cells (ECs), however, with age these predicted interactions shift. Cell-cell interactions of 2mo WT macrophages are equally distributed between a variety of ECs and fibroblasts (**Figure 3E**). However, at 24mo, WT macrophages interact most strongly with endocardial ECs. In contrast, in the aged STING^-/-^ hearts, macrophages appear to instead prioritize interactions with the lymphatic endothelial cells (LECs) and smooth muscle cells (SMCs) (**Figure 3E**). Monocytes from aged STING^-/-^ hearts also appear to interact strongly with LECs compared to WT monocytes, which instead split their interactions between non-lymphatic EC and fibroblasts (**Figure 3E**). Comparative analysis of macrophages and monocytes from STING^-/-^ livers reveals a specific loss of their interactions with stellate cells (fibroblasts), which is otherwise an interaction favored by aged WT macrophages (**Figure 3F**). Finally, like the heart and liver, STING regulates interactions between macrophages and the variety of epithelial cells found in the kidney, as in the absence of STING, macrophages favor interactions with fibroblasts in aged kidneys (**Figure 3G**). Collectively, these shifts in inflammatory immune signatures and cellular profiles uncovers important consequences for cell-cell interactions in aging tissues and in the absence of STING. Our data highlight how STING-signaling *fine tunes* the immune landscape of aging tissues: downregulating some genes and upregulating others in a highly cell-selective manner to impact cell-cell communication.

### STING, A MASTER REGULATOR OF IMMUNE RESPONSES, REINFORCES ORGANISMAL LONGEVITY AND HOMEOSTASIS

We next investigated the consequence for the unexpected role STING plays in inherent regulation of tissue immune landscape. Despite studies evaluating the role of STING in accelerated models of aging, and in models where treatment with antagonists is used to propose pathological roles for this pathway, the role of STING in physiologic and chronological aging has remained unknown ^3,6,24^. It is necessary to clarify, as aging itself is not a disease but rather a state of flux that may predispose one to chronic diseases, and our understanding of the immune basis for healthy vs. pathological remains biased towards the latter. To fill this outstanding gap, we first evaluated if aging impacts recirculating inflammatory cytokine/chemokine levels in otherwise unmanipulated STING-deficient (STING^-/-^) and age-matched wildtype (WT) mice (**Figure 4A**). Contrary to expectations, we note significantly increased levels of TNFα, IL-6, IL-12, IFNγ and GMCSF in STING^-/-^ sera (**Figure 4A**), suggesting a heightened inflammatory state in the absence of STING. Routine histopathology reveals greater numbers of infiltrating cells in the heart, liver, and kidneys of STING^-/-^ mice compared to WT controls (**Figure 4B-D**) and further correlates with increased systemic inflammation. Specifically, aged STING^-/-^ hearts have a larger number of infiltrates and increased numbers of cells around vasculature (**Figure 4B and S5A**) compared to age-matched WT. We also note increased cellular infiltrates in STING^-/-^ livers although many are interstitially localized and not proximal to the vasculature as in WT livers (**Figure 4C and S5B**). This altered localization pattern of infiltrates is in part complicated by the high degree of stromal disorganization notable in aged STING^-/-^ livers (**Figure 4C and S5B**). STING^-/-^ kidneys present with remarkable histopathological changes that are represented by increases in tubular infiltrates, size of Bowman’s capsules, and the thickness of parietal epithelial cell layer surrounding the Bowman’s capsules (**Figure 4D and S5C**). Overall, tissues from aged STING^-/-^ mice are substantially reorganized, with tissue stroma specifically appearing unstructured and disordered compared to similarly aged WT controls (**Figure 4B-D**). Importantly, these changes in tissue architecture result in more stochastic localization of infiltrating immune cells within STING^-/-^ tissues than WT controls. Not all these changes to tissue architecture may be dependent upon STING, especially if STING expression is restricted to key cell types in these tissues. To assess cell-specific expression patterns of STING (*Tmem173*), we leveraged our scRNAseq data and note that STING is broadly expressed, with preferential enrichment in the myeloid compartment and key stromal cells (**Figure S5D**). Thus, the expression of STING is likely a key driver of these changes in tissue composition with aging. Given these unexpected increases in markers of inflammation in STING^-/-^ mice, we next evaluated how STING-deficiency impacts lifespan. Consistent with the higher levels of frailty associated cytokines^25–27^ IL-6, TNFα and IL1β (**Figure 4A**), and the overall proinflammatory milieu in STING^-/-^ mice, we note that STING-deficiency significantly shortens lifespans (**Figure 4E**). Importantly, these changes were noted only in male STING^-/-^ mice, not females (**Figure S5E**), suggesting that STING may influence the sexually dimorphic nature of longevity. Collectively, these data completely upend current paradigms that place STING-signaling as a *driver* of chronic inflammation and shorter lifespans. Furthermore, our data underscore the high relevance of evaluating the role of molecules like STING in physiologic aging for accurate insights into their role in homeostasis.

**Figure 4:**
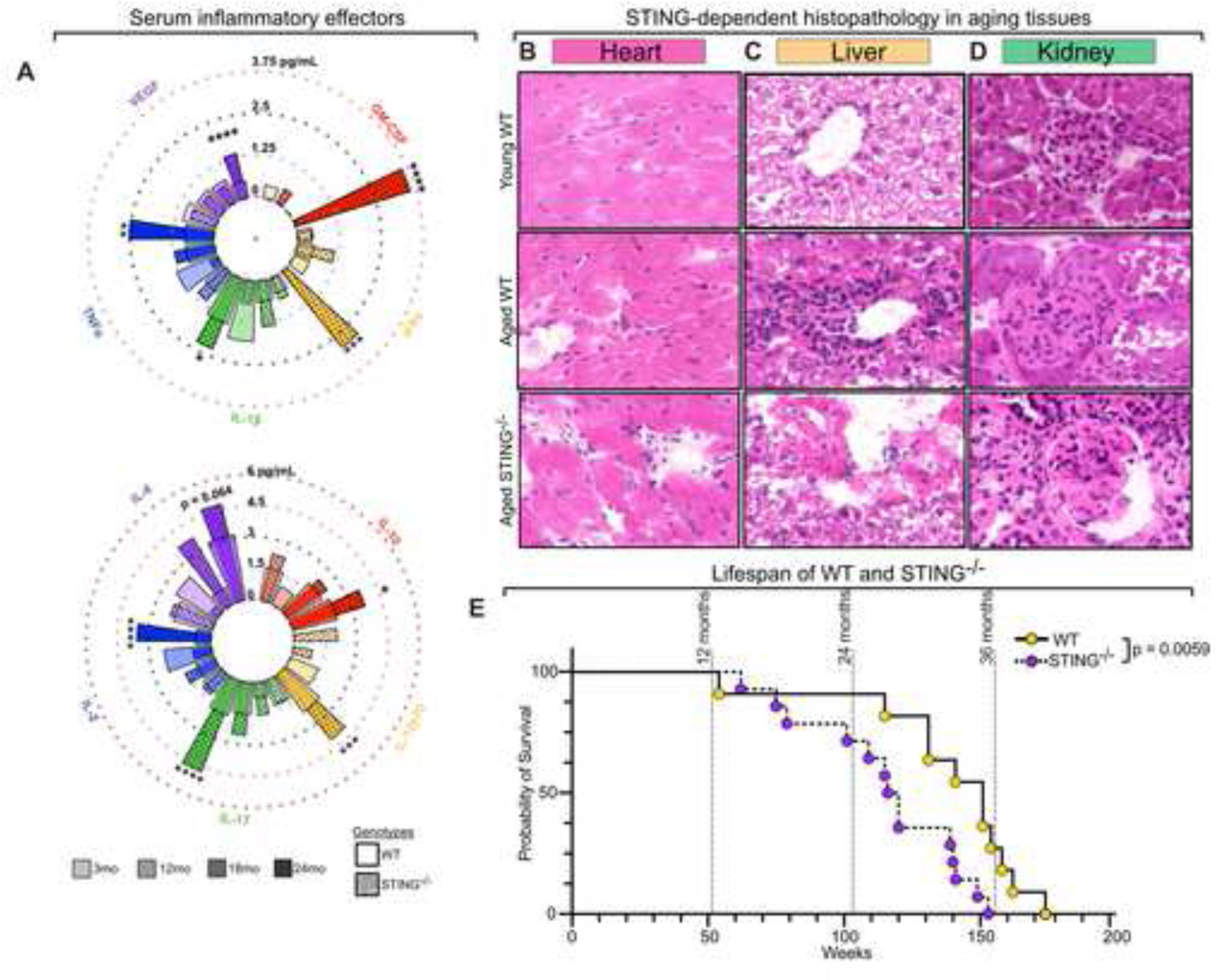
STING regulates systemic and histological inflammation to reinforce longevity. (A) Radial bar chart depicting serum cytokine and chemokine concentrations (pg/mL) from 3, 12, 18, and 24mo STING^-/-^ (filled, dotted bars) mice and WT (solid, transparent bars) measured by multiplex ELISA (N = 4-5). **(B-D)**. H&E analysis of young (2mo) WT and aged (26-32mo) WT and STING^-/-^ heart, liver, and kidney samples. **(E)** Kaplan-Meier survival probability curves comparing WT (solid line, yellow circles) and STING^-/-^ (dashed line, purple circles) male mice. Statistics were calculated with a one-way ANOVA with Bonferroni correction (A) or a Mantel-Cox Logrank test (E) \**p ≤ 0.05*; \*\**p ≤ 0.01*; \*\*\**p ≤ 0.001,* \*\*\*\**p ≤ 0.0001*.

### HOMEOSTATIC TISSUE-RESIDENT MACROPHAGE POPULATIONS ARE SIGNIFICANTLY ALTERED BY STING-SIGNALING

Tissue immune signatures in STING^-/-^ mice reveal significant bias towards increased macrophage activation. A role for STING in the immune responses of bone-marrow derived macrophages (BMMs) is recognized, however, its impact on a critical subset of macrophages – tissue-resident macrophages (TRMs), is completely unknown. TRMs have distinct, often mixed, ontogeny from BMMs, are finely tuned to their resident tissues, and are essential for key functions associated with tissue remodeling, development, and tissue homeostasis ^10,28–33^. Thus, to explain the essential dysregulation of tissue immune landscapes, we next sought to understand the physiological impact of STING-deficiency on TRMs. Subsetting and reclustering the macrophage populations in each tissue revealed significant heterogeneity, which has been reported by others (**Figure 5A-C**)^34^. The notably conserved TRM subclusters in each tissue were Folr2/Mrc1^+^ (cardiac), Tim4^+^ Kupffer cells (hepatic) and Mrc1^+^ (renal) TRMs (**Figure 5A-C**), closely resembling TRMs previously described to be long-term, homeostatic Tim4/Folr2/Lyve1 positive (TLF) TRMs (referred to as ‘TLF-like TRMs’)^34^. We identified other previously described TRM subsets such as *Ccr2*^+^ and *MhcII^hi^* TRMs in the three tissues (**Figure 5A-C**)^34^. In addition, we annotated several novel TRM subsets based on curated gene expression and pathway analysis (**Figure 5A-C, S6A-F).** Of these, a notable and novel TRM cluster common to all three tissues, expresses, among other key genes, the long non-coding (lnc)-RNAs *Malat1*, *Neat1* and/or *Xist*, which we annotate as *lncRNA*-TRMs (**Figure 5A-C, S6A-C)**. Stressed/inflammatory TRMs (characterized by inflammatory, senescence and/or unfolded protein response signatures) were also found in all three tissues (**Figure 5A-C, S6A-F).** The kidney and liver contain metabolically active TRMs, based on the expression of anabolic lipid and amino acid modifying networks (**Figure S6D-F**). Finally, we also identify unique populations of TRMs specific to each tissue, cycling (cell-cycle checkpoint signatures) TRMs in the liver, and pericardium associated (PC) TRMs in the heart (**Figure 5A-C**)^35^. Thus, our analysis captures previously characterized and novel subsets of TRMs that together describe a vital and plastic tissue-immune population.

**Figure 5:**
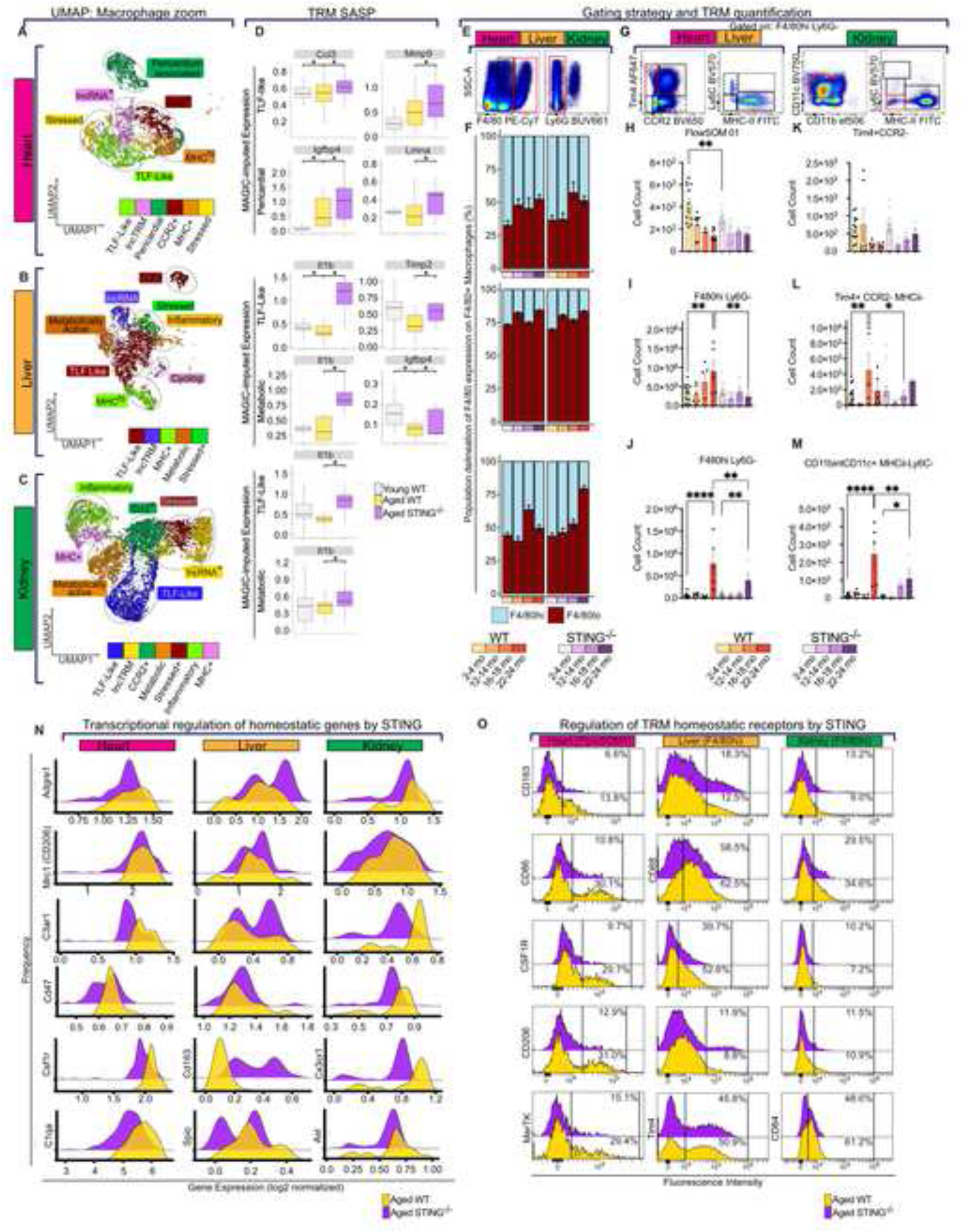
STING deficiency reshapes homeostatic TRM populations. (A-C) UMAP of subsetted and re-clustered macrophages from combined datasets of WT and STING^-/-^ hearts **(A)**, livers **(V)**, and kidneys **(C)**. **(D)** Boxplots denoting gene expression distributions of SASP- associated genes from annotated tissue resident macrophages (TRMs). Boxplots indicate upper and lower range, first and third quartile, and median. Grey, 2mo WT; gold, 24mo WT; purple, 24mo STING^-/-^. Bonferroni adjusted p-value thresholds are indicated as \**p ≤ 0.1.* **(E)** Flow cytometry gating strategy to identify macrophages from the heart, liver, and kidney. **(F)** Stacked bar plot of F4/80^hi^ (light blue) and F4/80^lo^ (dark red) macrophages from the heart, liver, and kidney. Ages and genotypes annotated below. **(G)** Continued from gating in **(E)**, flow cytometry gating strategy to identify resident macrophages in the heart, liver, and kidney. (H**-M)** Individual dots represent mice. Quantification of macrophages from the heart FlowSOM01 (aka TRM-1 or FlowSOM metacluster 1). Tukey correction adjusted p-value thresholds are indicated as \**p ≤ 0.05*;\*\**p ≤ 0.01*; \*\*\**p ≤ 0.001,* \*\*\*\**p ≤ 0.0001.* **(H)**; Tim4^+^ CCR2^-^ **(K)**; liver, F4/80^hi^ Ly6G^-^ **(I)**, Tim4^+^ CCR2^-^ MHCii^-^ **(L)**; and kidney (F4/80^hi^ Ly6G^-^ **(J)**, CD11b^int^ CD11c^+^ MHCii^-^ Ly6C^-^ **(M)**. Ages and genotypes as indicated in **(F)**. **(N)** scRNAseq based "ridgeplots" of gene expression for homeostasis related genes critical for scavenging and efferocytosis functions of TLF-like TRMs, from the heart, liver, and kidney. Genotypes annotated as in (d); WT; gold, STING^-/-^ purple. **(O)** As in (n), histograms of homeostatic surface receptors on TRM-1 (heart) and F4/80^hi^ macrophages (liver, kidney). 24mo samples are concatenated into representative images (n = 5 WT, 4, STING^-/--^ ). Percent positive cells are annotated as shown. Genotypes annotated as in (d, n); WT; gold, STING^-/-^; purple. Statistics calculated using a Wilcoxon Rank Sum test (d) or One Way ANOVA (h-m).

A cell intrinsic role for STING in TRM subsets is further supported by a relative enrichment of STING (*Tmem173*) in multiple TRMs with aging (**Figure S7A-C**). Using known markers of long-term residency (e.g. Tim4, Folr2, Lyve1, Mrc1), we were able to identify 2 main clusters in each tissue as likely being long-lived TRMs, namely the Folr2/Mrc1^+^ and PC-associated (cardiac), Tim4^+^ and metabolic (hepatic) and Mrc1+ and metabolic (renal) TRMs (**Figure 5A-C, S6A-C**). Differential gene expression (DGE) analysis of the metagene signatures in TRMs reveal that STING-deficiency does not impact INF and ISG genes significantly in the long-term TRMs (**Figure S7D-E).** Indeed, at 24mo of age, the majority of the INF genes in these large Mrc1^+^ or Tim4^+^ subsets are shared between WT and STING^-/-^ TRMs with some notable differences such as the expression of *Cxcl13, Ccl8, Cd14, Ccl6* and *Csf1r* in STING^-/-^ TRMs **(Figure S7D**). Although the loss of STING does not appear to alter the induction of senescence based on *Cdkn1a*/*2a* expression in macrophages (**Figure S7F-H**), it paradoxically correlates with *greater* expression of known inflammatory SASP genes *Mmp9, Il1b, Lmna, Timp2* and *Cxcl1* in long-term TRMs (**Figure 5D**). Aging has been shown to reorganize the TRM pool and, given the shifts in senescence gene expression, we wondered if STING-deficiency further impacts TRM population size during aging. To quantitatively address this, we utilized high-dimensional, spectral flow cytometry (26-33-marker). We selected tissue associated macrophages by their relative expression of F4/80 (i.e., F4/80^hi^ and F4/80^lo^) which defines the two largest classes of tissue-associated macrophage populations (**Figure 5E-F**). F4/80^lo^ macrophages predominate tissues with aging but were similarly distributed between WT and STING^-/-^ tissues (**Figure 5F**). F480^hi^ macrophages are often considered synonymous with long-term TRMs, which are regarded the gatekeepers of tissue integrity, however recent analyses have revealed that historical classification of TRMs as a homogenous subset, may be highly reductive ^10,11,30,32,34,36–38^. Thus, we selected the tissue-associated F4/80^hi^ macrophages as a parent population to generate un-biased Self-Organizing Maps (Flow-SOMs). These Flow-SOM “metaclusters” were used as a basis to further categorize TRM subsets and identified 7 TRM populations with phenotypic marker differences in each tissue (**Figure S8A- C**). Overlaying the Flow-SOM metaclusters on a UMAP generated from F4/80^hi^ cells reveals these heterogeneous clusters of TRMs in the three tissues (**Figure S8D-F)**. In the heart, TRM metacluster-1 (TRM-1) was significantly abrogated as early as 3mo of age in STING^-/-^ mice (**Figure 5H and S8D**), suggesting that STING is involved in the early reorganization of a large proportion of the cardiac resident macrophage population. In contrast, the STING^-/-^ liver and kidneys, we see a profound decrease in multiple subsets of F4/80^hi^ TRMs (**Figure 5I-J and S8C-D**), suggesting that STING expression regulates the maintenance of significant populations of resident macrophages in these tissues.

To directly correlate these findings with our scRNAseq analysis, we specifically evaluated previously characterized long-term, resident TRMs in WT and STING^-/-^ tissues using defined gating strategies that rely on Tim4 expression in the heart and liver and CD11c in the kidney (**Figure 5G, K-M)**^34,39,40^. In the heart, bulk Tim4^+^ cardiac TRMs are not significantly altered, but trend lower in STING^-/-^ mice by 12mo of age. (**Figure 5K**). Alternatively, in the aging liver and kidneys, STING^-/-^ tissues have significantly lower numbers of Tim4^+^ and CD11c^+^ TRMs at 18mo and 24mo respectively (**Figure 5L-M**). Collectively, our data reveal that with aging, long-term TRM populations are smaller with STING-deficiency but reveal a tissue-specific chronology of TRM loss.

Given the dramatic reduction in TRM subsets in STING^-/-^ tissues, we next evaluated if markers of routine homeostatic functions, such as receptors for scavenging and phagocytosis were impacted. The subsets selected for DGE analysis form a large percentage of the TRM populations, highly suggestive of key roles in tissue maintenance. A careful evaluation of a new metagene signature for homeostatic maintenance (**Supplemental Information 1**) in these populations suggest that indeed, homeostatic signatures are altered with STING-deficiency in TRMs (**Figure S8G-I)**. Many homeostatic genes responsible for scavenging, phagocytosis, and core metabolic function appear to be impaired in STING^-/-^ TRM subsets and include *Adgre1, Cd47, C3ar1, Axl, Mrc1, Cx3cr1, C1qa, Csf1r, Cd163, Spic* and others (**Figure 5N)**. We corroborated this transcriptional data by assessing the levels of key homeostatic cell-surface markers such as CD163, CD206 (Mrc1), Tim4, MerTK and others using spectral flow cytometry (**Figure 5O**). Importantly, CD163, Csf1r, CD68, CD64, CD206 and Tim4 are all downregulated on F4/80^hi^ macrophages with aging in the STING^-/-^ tissues (**Figure 5O**), suggesting that the STING-deficiency has a critical impact on homeostatic gene and protein expression in TRMs in addition to regulating the size of the TRM pool.

### THE STING-PATHWAY ENCODES VIABILITY AND VITALITY OF TRMS

Several mechanisms could explain the loss of TRMs in STING^-/-^ tissues. To understand how STING signaling shapes the landscape of tissue-resident macrophages and monocytes, we used GSVA analysis to identify relative enrichment of GO: Biological Processes in aged TRMs. Comparing the magnitude of change in aged WT and STING-/- TRMs we identified a number of pathways that negatively regulate cell death processes, specifically sensitivity to cell-death via oxidative stress, are commonly dysregulated in STING-/- TRMs regardless of tissue of origin (**Figure 6A**). These data indicate that STING-deficiency may leave key subsets of TRMs more prone to cell-death. Indeed, confocal imaging and analysis reveals that aged tissues like the liver accumulate more dying TRMs (as identified with TUNEL staining), *in situ* (**Figure 6B-C**). Importantly, the numbers of TUNEL+ TRMs are significantly increased in the absence of STING in the liver (**Figure 6C**). We then sought to evaluate if, in fact, STING^-/-^ TRMs are more susceptible to death by common (apoptotic, pyroptotic, oxidative or necroptotic) stressors. We used unelicited, resting large peritoneal macrophages, previously shown to be the long-term TRM in the peritoneum ^38,41^, and BMMs as TRM and monocyte-derived macrophage populations respectively, and probed their ability to respond to death stimuli (**Figure 6D**). Indeed, STING^-/-^ TRMs show accelerated cell-death kinetics compared to WT TRMs (**Figure 6D**). Overall, BMMs from either genotype behave identically, underscoring the idea that tissue-specific programming plays a key role in specifying the sensitivity of macrophages to cell-death stimuli. Specifically, in agreement with GSVA analysis, we noted increased rates of cell-death in STING^-/-^ TRMs in response to the DNA-damaging agent, etoposide (apoptosis) and hydrogen peroxide (oxidative stress) (**Figure 6D**). Overall, these novel findings emphasize the critical role STING plays in tuning the responsiveness of TRMs to excessive DNA-damage and increased oxidative stress, both highly prevalent in aging tissues^42–45^.

**Figure 6:**
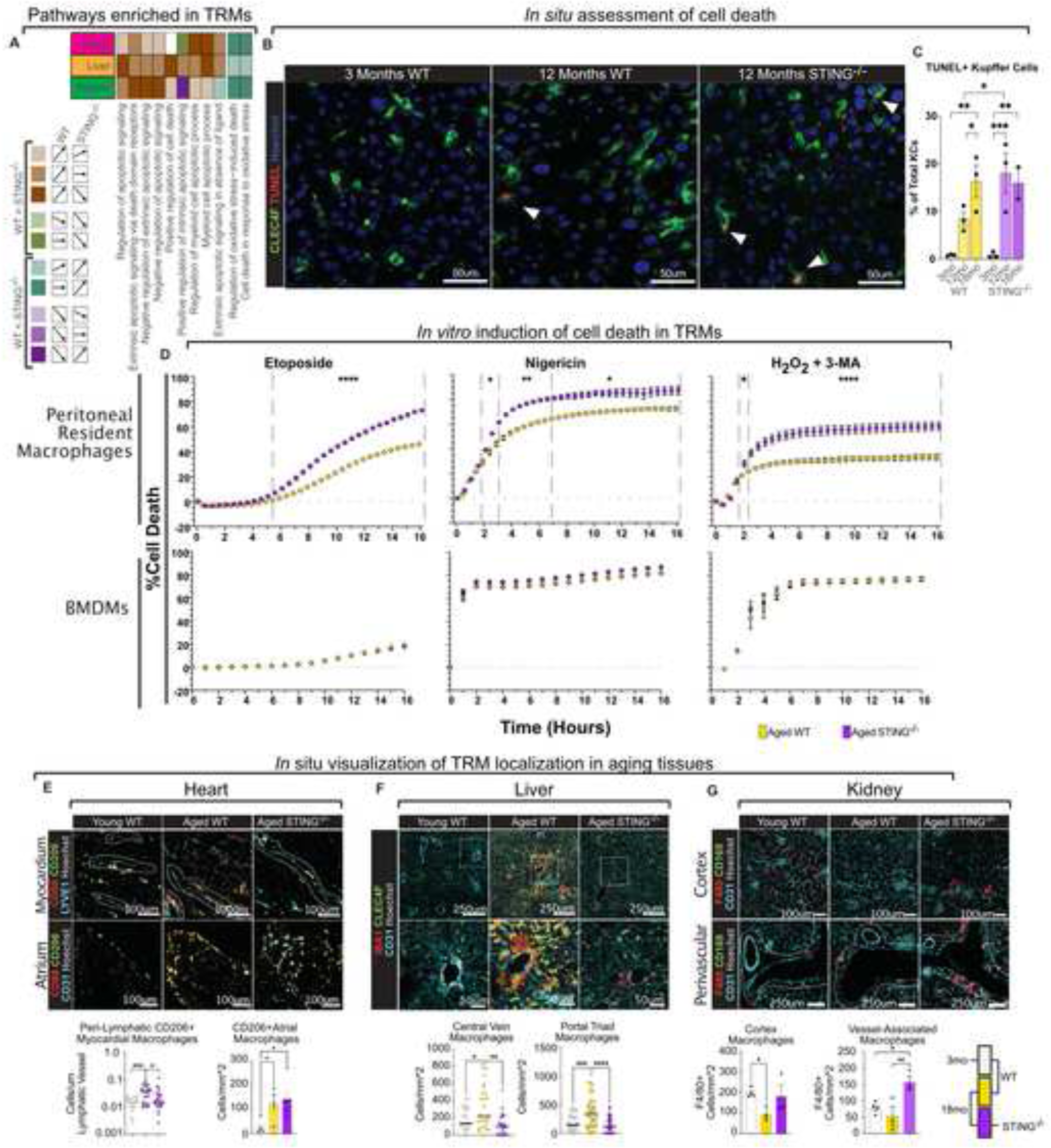
TRM vitality and tissue distribution require the STING pathway. (A) Summary of GSVA enrichment scores of GO pathways related to cell death in TLF-like resident macrophages. Shading and colors indicate trends of relative pathway scores in 24mo WT and STING^-/-^ relative to 2mo control TLF-like macrophages at a significance threshold with an FDR adjusted p-value < 0.05. **(B)** Immunofluorescent TUNEL staining of CLEC4F+ Kupffer cells (liver resident macrophages). White arrows indicate dual TUNEL and CLEC4F staining indicating dying Kupffer cells. (Red, TUNEL; green, CLEC4F; blue, Hoescht. 3 fields of view per tissue; number of independent tissues, N = 3) **(C)** Quantification of TUNEL+ Kupffer cells as percent of total CLEC4F^+^ Kupffer cells (right). **(D)**. In vitro cell death assays testing vitality of 16-week resident macrophages (large peritoneal macrophages) and bone marrow derived macrophages (BMDMs) by administration of Etoposide (left), Nigericin (middle), and H2O2 + 3- MA (right). Data points are average of n=5 technical replicates. Figure representative of N = 4 biological replicates. **(E-G)** Visualization and quantification of TRM subsets in relation to local structures and vasculature. At least two fields of view per liver, kidney, or entire atria (N = 3-5). **(E)** CD68^+^, CD206^+^ macrophages and CD31^+^ endothelial cells in the atrium and LYVE1^+^ lymphatic venules in the myocardium from 3mo and 18mo WT and STING^-/-^ heart. Peri- lymphatic localization of CD68^+^ and CD206^+^ macrophages are quantified as cells / length lymphatic venule in µm. (CD68, red; CD206, green; CD31/LYVE1, cyan; Hoescht, white). **(F)** IBA1^+^ and CLEC4F^+^ macrophages/Kupffer cells and CD31^+^ endothelial cells from WT and STING^-/-^ liver. Kupffer cell polarization around the central vein is quantified below. White dashed circles indicate central vein. White dashed boxes indicate regions for greater magnification. (CV, central vein; PT, portal triad. (IBA1, red; CLEC4F, green; CDE31, cyan; Hoescht, white). **(G)** F4/80^+^ and CD169^+^ macrophages and CD31^+^ endothelial cells from WT and STING^-/-^ kidneys. White dashed areas indicate perivascular space. Density of macrophages in the renal cortex and vascular polarization of macrophages are quantified. (Young, 3mo; Aged 18mo. F4/80, red; CD169, green; CD31, cyan; Hoescht, white). Statistics calculated using One Way ANOVA (d, e-g) or Two Way ANOVA (c) \**p ≤ 0.05*; \*\**p ≤ 0.01*; \*\*\**p ≤ 0.001*.

The changes in numbers of TRMs may result in a redistribution of TRMs in STING^-/-^ tissues, thus, we looked closely at the organization of TRMs in these tissues by confocal microscopy and immunofluorescence (**Figure 6E-G**). Importantly, we note that in the heart, CD206^+^ TRMs tend to accumulate in peri-lymphatic areas in WT and their homing here has been noted to be important for maintenance and appropriate functioning of the cardiac lymphatics^46–50^. However, this association is significantly decreased in the STING^-/-^ hearts (**Figure 6E**). In the liver, the zonation of TRMs outside the central vein (CV) boundaries and around portal triads (PT) is particularly critical in maintaining optimal liver function ^36,51,52^. In aging livers, Clec4F^+^ Kupffer cells (KCs) migrate and cluster around the CVs, revealing perturbations in their localization (**Figure 6F**). Intriguingly, Clec4F^+^ KCs are dramatically reduced in STING^-/-^ livers from mice and non-macrophage immune cells are instead found clustering around the CV with observable and significant disorganization of the stroma (**Figure 6F**). Furthermore, PT associated TRMs are also lost in STING^-/-^ livers. Similarly, in aged kidneys, the overall decrease in F480^+^CD169^+^ TRMs is notable in both WT and STING^-/-^ however F4/80^+^CD169^-^ macrophages are specifically and significantly only associated with the medullary vasculature in STING^-/-^. This shift in macrophage phenotype and re-localization (indicating infiltrating macrophages) proximal to vasculature in the kidneys (**Figure 6G**) is reminiscent of changes noted in some renal diseases ^53^. Altogether, these changes in STING^-/-^ TRMs suggest that STING-signaling specifies TRM tissue numbers and distribution by regulating their responsiveness to cell-death cues.

## DISCUSSION

Here we reveal that tissue-autonomous immune fingerprints are the intrinsic landscape upon which cell:cell interactions and age associated disruption of tissue architecture rest. Stromal and myeloid cells drive the component inflammatory, interferon-inducible and senescence gene profiles to shape the composite immune fingerprint of aging tissues. We uncover the paradigm-shifting, mechanistic contribution of the ER-resident innate-immune protein, STING, in shaping these immune fingerprints during physiologic aging. We reveal that STING *represses* both systemic and tissue inflammation, thereby rendering it critical for organismal longevity. Mechanistically, we identify the pan-tissue dysregulation of key homeostatic cell types, tissue-resident macrophages, in the absence of STING. The contraction of TRM populations in STING-deficient tissues is linked to their loss in expression of homeostatic genes and increased expression of key inflammatory markers in the tissues. Furthermore, STING-deficiency in TRMs specifically renders them, but not monocyte derived macrophages, more susceptible to cell-death triggers. STING-deficient TRMs are particularly sensitive to cell-death induction by DNA-damage and oxidative stress, both widespread features in aging tissues. These data implicate a role for STING-signaling in modulating the TRM response to hallmark age-related changes in tissues. Finally, we reveal that these TRM-intrinsic and extrinsic roles for STING also result in TRMs being consigned to unusual tissue sites, concomitant with an overall striking lack of stromal organization in STING-deficient tissues with age. Indeed, preserving STING-signaling long-term has beneficial effects on health-spans and lifespans during aging.

A comprehensive characterization of tissue inflammation with age, together with the detailed knowledge of the cellular players, molecular pathways and their interactions are particularly relevant for studies on lifespan extension, a sought-after ‘anti-ageing’ effect^54,55^. Our study tackles each premise of this assertion and reveals first, that INF, ISG and SEN signatures mark young *and* aging tissues with highly unique immune fingerprints. Second, overlaying the immune fingerprints upon tissue-associated cell types surprisingly reveals that, INF, ISG and SEN genes are not each expressed by distinct cell subsets. Contrary to expectations, expression of all three metagenes relies on critical subsets of stromal and myeloid cells, that coordinate the simultaneous expression of the three signatures. We also reveal that these tissue-based immune signatures are not merely derivative of recirculating serum cytokine/chemokine signatures but represent substantial immune signaling in tissue-autonomous cell-types. This analysis highlights, for the first time, the importance and diversity of the tissue-autonomous immune responses and reveals that the composite tissue signature is a combination of immune gene *parcels* from key cellular compartments. That these unique immune parcels shift with aging and with STING-deficiency emphasize that the *identity* of the metagenes expressed in various cell subsets rather than the expression levels of specific genes, is a critical determinant of tissue health.

The identification of a molecular pathway that may impact one if not multiple aspects of these tissue-based changes was a primary goal in our study. The cGAS/STING pathway has been described in several settings as a critical determinant of pathological signaling^56^, making STING, with its capacity to bind small cyclic molecules, a very attractive pharmaceutical target. In the context of aging, studies have suggested that abrogating STING-signaling may ameliorate senescence associated pathological outcomes in cells like fibroblasts^6,24,56–58^. A caveat of these studies is the fact senescence measurements have been universally low, with only 7-10% of the cell population being positive for senescence markers like *Cdkn2a* (p16) even in advanced aging^59,60^. These low numbers may at least be explained by the inconsistent expression of markers of senescence and have necessitated the use of more comprehensive gene-sets to identify senescent cells^61^. A further limitation of studies that implicate cGAS/STING in aging is the use of model systems that result in inadvertent accumulation of cytosolic DNA, such as seen with the YAP/TAZ, SIRT6 or the mitochondrial polymerase (*Polg*) deficiency^4–6^. Thus, studies implicating a role for STING in pathological aging, primarily through senescence or accumulation of cytosolic DNA due to mutations in core cellular machinery, may not fully capture its role in physiologic age-associated changes.

A recent study by Gulen et. al. reveals that treating aging animals short-term with a STING-antagonist (H- 151) reverses the expression of inflammatory immune signatures classically known to be STING-dependent.^24^ Moreover, they show that these same genes are altered in tissues like the brain, kidneys and liver from *Sting1*^-/-^ mice^24^. A caveat from this study is that while they address aspects of how STING- antagonist treatments may ameliorate age dependent immune gene expression in the brain or kidneys, their long-term impact on lifespans and overall health is not measured. This is a gap of critical relevance, as any treatment with senolytics and/or STING antagonists approved for aging individuals, will be expected to be prescribed as a life-long treatment.

Furthermore, at least 4 STING LOF polymorphic alleles are well known and stratify in the human populations at frequencies of >50%^8,9,62^. Importantly, the strongest LOF alleles are found in ethnic minorities, historically known to suffer the most from age-related complications^9^. These data suggest that the correlation between loss of STING function and aging needs to be defined in a longitudinal analysis of aging associated changes. Indeed, humans with these LOF alleles live lifelong with these polymorphisms, it is critical to assess how deficiency of STING at birth can reshape immune landscapes and reframe aging-associated decline. A categorical assessment of STING in lifespans and physiological age-associated systemic inflammation is lacking, we thus began by evaluating exactly these parameters in STING-deficient mice. We find that STING plays roles in *suppressing* pathological aging and promoting lifespans, that have not been previously described and upend its proposed paradigmatic pathological mechanisms in aging. STING-deficient mice have shorter lifespans, increased inflammatory serum cytokine burden and significant histopathological changes compared to littermate WT mice. It is imperative to note here that the changes in lifespans are significant only in *male* STING-deficient mice, as female mice do not have significantly different lifespans compared to their WT littermates. This sexually dimorphic role for STING in extending the lifespan of males but not females is intriguing, other parameters are identical between males and females. Moreover, *ex vivo* measurements such as cell-death were not impacted by the sex of the mouse. Our study marks a separate description of how male mice deficient in STING fare worse in chronic disease and aging, as STING-deficiency accelerated disease in male mice of a model of SLE and also bone loss with aging^63–65^. We speculate that this STING-dependent difference is in part driven by the hormonal milieu and remains a key point of inquiry in our future studies.

We speculate that STING-dependent histopathological changes and serum cytokine levels likely reflect critical changes in tissue immune signatures. This is indeed the case, as STING-deficiency leads to striking changes in the tissue-immune fingerprints comprised of INF, ISG and SEN genes. What is perhaps most remarkable is that the loss of STING does not merely abrogate expression of genes upregulated in aged WT tissues, but instead leads to a recalibration of the tissue immune signature. In many scenarios this is reflected by an upregulation of proinflammatory genes like *Cd14*, *Pycard, Ptgs2* and *Il1b* amongst others in aging STING-deficient tissues. The increased expression of inflammatory genes in STING^-/-^ tissues may indicate an increased predisposition to inflammation in the presence of the right triggers or ongoing inflammatory events driven by common danger-associated molecular patterns (DAMPs) found in aging tissues. The upregulation of classical inflammatory genes like *Il1b*, *Ptgs2*, and others suggest that STING-signaling suppresses the *ad hoc* expression of these genes by unknown pathways. As stromal cells and resident myeloid cells express key portions of these gene sets, it is also unknown if the mechanism by which STING suppresses gene expression is similar in the distinct cell types. Thus, our studies have identified the existence of uncharacterized downstream mechanism(s) by which STING regulates the expression of key immune response genes in multiple tissues.

One mechanism by which STING may modulate these far-reaching consequences in tissue immunology is through the common regulation of cellular elements key to tissue function and homeostasis. Stromal elements are unique to each tissue. Although STING clearly plays a role in shaping their interactions with other cells and overall histological organization, stromal cells do not represent a common cell-type that may unify the multi-tissue and systemic effects of STING-deficiency. This explains our interest in the highly conserved, pan-tissue, homeostatic cell type – TRMs, as a potential critical cellular player ^31^. TRMs are an ontologically heterogeneous macrophage subgroup, highly attuned to their microenvironment to facilitate imprinting, remodeling and other core tissue functions^11,31,33^. Our understanding of the biology of resident macrophages has recently exploded, with the advent of lineage tracers and single-cell technologies^23,32,38,66,67^. While core transcription factors such as *Gata6*, *Id2*, and others have been proposed to determine the lineage specificity of TRMs in specific tissues^11,32,33^, no known role for innate immune pathways in specifying TRM populations or homeostatic properties has been forthcoming. It was thus, doubly surprising that not only does STING-signaling support the size of TRM populations, but also regulates the expression of homeostatic cell-surface markers. Decreases in TRM numbers are associated with poorer prognosis in multiple chronic disorders^37,68–70^. Thus, the early and sustained lowered numbers in TRMs in STING^-/-^ tissues suggests a highly critical role for this pathway in specifying TRM fate with age-induced changes in tissue biology. Indeed, as TRMs are carefully paired with stromal cells in many of the tissues^38,66,71^, an intrinsic change in TRM biology is likely to directly impact these multi-faceted interactions with other cell types in the tissue. Indeed, we see some indications of these changes with our initial RNA-magnet experiments, where we note that STING-signaling shapes the interaction of macrophages with key stromal anchors in the heart (LECs), liver (stellate cells) and kidneys (fibroblasts). Intriguingly, we note similar changes with direct immunofluorescence, where in the heart, STING^-/-^ TRMs lose their apparent association with Lyve1^+^ ECs (considered overlapping with LECs). In our hands many of the Lyve1^+^ EC associated TRMs were also Lyve1^+^, the loss of this interaction suggests that vascular integrity may be impaired and drive fibrosis as reported previously^29,47,50^. Similarly, in the liver, we see a significant de-zonation of STING^-/-^ TRMs. The loss of TRMs in these specified niches is likely to predispose the tissue to increased infiltrates, fibrosis and pathogen susceptibility^36,51,52,72,73^. Moreover, the loss of interaction with stellate cells in STING^-/-^ livers may have another outcome, liver fibrosis is a common feature of aging livers and considered pathological^72,74–78^. However, fibrosis may have the inadvertent benefit of maintaining structural integrity of the tissue architecture. Indeed, STING^-/-^ livers appear highly disorganized compared to age-matched WT, suggesting some inherent loss of architectural hierarchy. In contrast, in the STING^-/-^ kidneys, the increased interaction with fibroblasts may disrupt kidney organization by increasing the incidence of fibrosis in these tissues. It is critical to note that while TRM loss is a common feature of STING-deficiency in these tissues, the effect of TRM loss in these tissues has many trajectories. Overall, a domino-like effect is likely to ensue if TRM-behavior is compromised, leading to the lack of organization and failure of cell:cell cues to maintain order. However, it remains to be ascertained if a TRM-intrinsic role for STING is sufficient for safeguarding against age-associated shifts in tissue pathology and the overall shift in lifespans.

Finally, we explore how STING-deficiency modulates TRM cell numbers in tissues. Changes in TRM populations impact several tissues with aging ^34,52,79^. Yet, the basis of this population change is unknown, and several overlapping mechanisms may explain it: a) Cell-death; b) Egress from the tissue; c) Infiltration of monocyte-derived macrophages, etc. d) Variations in proliferative cues. Here, we reveal that in the case of STING-deficiency, the contraction of TRM populations is in part due to increased sensitivity of STING^-^

^/-^ TRMs to key cell-death triggers. This is also essentially true of TRMs compared to monocyte-derived macrophages (BMDMs), where they reveal heightened sensitivity to cell-death triggers of a certain kind. Curiously, the key triggers to which STING^-/-^ TRMs appear most sensitive include etoposide and H2O2, these roughly correspond to triggers for DNA-damaged induced cell-death and cell-death in response to oxidative stress. Intriguingly, STING has previously shown to be critical in the immune response to genotoxic stress (chromosomal breaks, micronuclei, double-stranded DNA breaks, etc.)^56,80^. Here we reveal that STING also functions to commit a cell to survival by engaging this immune response and in its absence, the cell engages death pathways to cope with irreversible and continuing DNA-damage. The relationship between oxidative stress and STING is poorly characterized, although oxidation of STING is thought to inactivate its signaling functions and downstream immune response^81^. Here we show that increased oxidative stress in the absence of STING compromises cell viability and sensitizes TRMs to cell-death. It remains to be seen if this feature of STING (balancing immune response with commitment to cell-death) is specific for TRMs or can be generalized to other cell types.

Overall, our study focuses on a much-debated mechanism of aging by assessing STING-deficient mice that age chronologically and physiologically at various timepoints for age-associated changes. Thus, in these core aspects, our study addresses fundamentally different mechanistic contributions that STING signaling makes in age-associated decline and does so with relevance to lifespan analysis, comprehensive immune signature mapping and histological evaluation. Recent years have seen the rapid development and clinical testing of both therapeutics that target the cGAS/STING pathway for use during cancer, autoimmunity, infections and other immune conditions^56^. With new roles ascribed to the cGAS/STING pathway in senescence, these therapeutics are likely to be employed alongside existing senolytics with a goal to improve health-span and lifespan of elderly individuals. However, our studies emphasize the unanticipated impact of using cell-blinded, small-molecule therapeutics like cGAS/STING antagonists in aging individuals over the course of their lifespans. The prevalence of LOF polymorphisms in STING within the human population, necessitate a careful examination of how STING-antagonists interface with existing STING alleles, which has thus far been ignored. We believe that short-term treatment with senolytics and STING-antagonists may in fact have beneficial effects, especially in humans with no LOF in STING. However, our data now suggest that functional status of STING may play a critical role in stipulating drug-dosing for maximal benefit and minimal harm. The urgent need for comprehensive studies evaluating the linkage between STING-alleles and longevity or health is underscored by our findings. Furthermore, we reveal how, in the context of STING, immuno-protective pathways are interwoven with pathological ones and provide insight into why *brute force* treatments are often not successful in many disorders. Our study has unlocked several new research frontiers, with provocative consequences for multiple chronic disorders and aging associated dysfunction.

## ACKNOWLEDGEMENTS

We would like to thank Pilar Alcaide, Sasha Smolgovsky, and Zoie Magri for providing resources, reagents, and experimental support. We are grateful to the Tufts University Core Facilities for providing essential services (Flow Cytometry, Genomics), the Tufts University Comparative Medicine Services (DLAM, RBS) for animal support, maintenance, and husbandry. Support for this research was provided by research grants through the NIH (SS: R01AI142005; AP: R01AI167245), and through private companies/research foundations (SS: Arthritis National Research Foundation, Arthritis and Aging Research Grant, and a 10X Genomics Mini Grant).

## AUTHOR CONTRIBUTIONS

Conceptualization, SS, JWH; Methodology, JWH, KBS; Investigation, JWH, KBS, MS, KAC, RDB, ERR, AT; Writing – Original Draft, SS, JWH, KBS; Writing – Review & Editing, SS, JWH, KBS, SCB, MS, KAC, ERR; Funding Acquisition, SS; Resources, AP, AT; Supervision, SS.

## DECLARATION OF INTERESTS

The authors declare no competing interests.

## Methods

### Mice

Male and female wild type (WT, C57BL/6 background) and STING^-/-^ (Tmem173^-/-^, C57BL/6 background) mice were used for all aging and in vitro experiments. Aged experiments were organized into 3-months-old (mo) WT mice (Tufts Medical School), 12-mo WT mice (Tufts Medical School or National Institute of Aging (NIA)), 18-mo WT mice (Tufts Medical School or NIA), 24-mo WT mice (Tufts Medical School or NIA). Mice from the NIA were habituated for at least a month in the animal facility before experiments. Randomized groups of age-matched mice were also set up to determine longevity while Tufts University Division of Laboratory Animal Management (DLAM) performed daily health checks on each cage. STING^-/-^ mice (G. Barber, U Miami) were set up to match WT ages: 3-, 12-, 18-, 24-mo experimental groups (Tufts Medical School). All mouse strains were bred and maintained by Tufts Rodent Breeding Services (RBS) in accordance with the Institutional Animal Care and Use Committee (IACUC) and guidelines set by Tufts Medical School. Mice were housed under specific pathogen-free conditions in a 12-hour light-dark cycle. Mice were randomly allocated to aged groups and maintained with regular chow and water ad libitum unless euthanasia was required prior to the designated age per Tufts DLAM recommendations and requirements.

### Tissue collection and processing

Mice were euthanized by CO2 administration (4.5L/min) per IACUC regulations. After euthanasia blood was collected through cardiac puncture (25G) and allowed to clot at room temperature (RT) in RNase free 1.5mL Eppendorf tubes. Peripheral blood was spun in a refrigerated centrifuge at 10,000rpm for 10min at which point the serum was aliquoted and stored at -80°. Organs were harvested by blunt dissection into DMEM then embedded, and flash frozen in optimum cutting temperature cryo-embedding medium (OCT) and stored at -80° or dissociated to single cell suspensions for further processing for flow cytometry, *in vitro* analyses, or single cell RNA sequencing. Specifically, heart, liver, and kidney were minced with a razor blade and incubated with 200U/mL collagenase II and DNase I in serum free DMEM at 37°C with intermittent shaking (hearts; 20min, liver; 40min, kidney; 45min). After enzymatic digestions, suspensions were mechanically dissociated through a 70μm filter and red blood cells were lysed using ACK red blood cell lysis buffer per manufacturer’s instructions. Dead cells and other cellular debris were removed using the Miltenyi Debris Removal Kit following standard protocol. Cells were counted (Countessa, Thermo, with Trypan Blue viability stain) and aliquoted for experimental use.

### Flow cytometry

All steps were performed on ice or incubated at 4°C. After generating single cell suspensions as described above, cells were washed with Phosphate Buffered Saline (PBS, Corning) and resuspended in 100μl of serum-free PBS with Zombie NIR (ZombieNIR anti-mouse Live/Dead) for 15-30 minutes. Cells were washed (5min, 400rpm 2x with 2mL FACS buffer (2% FBS in PBS)) and resuspend in 50μl of PBS with FC block (anti-mouse CD16/32; clone 2.4G2) for 15 minutes, and then incubated with a master mix of antibody cocktail for 30 minutes (total of 100μl/ 2x10^6^ cells). Cells were then washed and centrifuged for analysis on the Cytek Aurora spectral flow cytometer (Cytek; Tufts Medical School). Single color controls were used on cells or beads depending on the prevalence of the marker on target cell types. Unmixing was conducted using SpectroFlo (Cytek) and data was analyzed with Omiq (Dotmatics).

### Fluorescence-Activated Cell Sorting (FACS)

For fluorescence-activated cell sorting surface staining was performed as described. Cells were stained with Zombie viability dye and surface markers for anti-mouse CD45 (30-F11), anti-mouse Ter119 (TER- 119), anti-mouse CD3(17A2/145-2C11), and anti-mouse CD19(6D5). CD45^+^ cells were sorted with the Legacy MoFlo (Cytomation; Tufts Medical School) into 1% BSA-PBS in 1.5mL Eppendorf tubes.

Samples were used for Chromium Single Cell Gene Expression sequencing analysis.

### Cryosectioning

Cryosections were generated as described above, and frozen tissues were sectioned at 5-15μm thickness with a Thermo Shandon Cryotome FSE to be mounted, air-dried, and stored at -80°C until hematoxylin and eosin (H&E) staining by iHisto INC (µm) or immunofluorescence (15µm). H&E staining was performed

per iHisto protocols and image acquisition was performed on Lionheart FX Automated Microscope (Tufts University).

### Quantification of resident macrophages in frozen sections

#### Imaging

Prior to staining slides were allowed to thaw to room temperature when OCT was removed with PBS and tissues were fixed with cold 4% paraformaldehyde (PFA) for 20 minutes and washed (5 minutes in PBS, 2x). Fixed tissues were blocked with 5% donkey serum (Thermo) in PBS and permeabilized with 0.3% Triton-X (Sigma) for 1 hour at room temperature (RT) before overnight incubation with primary antibodies diluted in blocking solution at 4^°^C. Following primary staining, tissues were washed before incubating with secondary antibodies diluted in blocking serum for 2 hours at RT. Finally, tissues were washed and incubated with Hoechst (Thermo) 1:2000 in PBS. Coverslips (Corning) were mounted using Fluoromount-G (Southern Biotech) and sealed.

#### TUNEL Staining

Tissue sections were processed through fixation as above, and then incubated with TUNEL (Roche) according to manufacturer’s instructions before proceeding to blocking.

#### Imaging

Immunofluorescent images were acquired using a Leica SP8 confocal microscope (43x or 63x objective) and analyzed using FIJI (v2.3.0) or CellProfiler (v4.2.1). At least 2 fields of view were acquired from at least 3 biological replicates for heart (myocardium, 0.8mm^2^), liver (0.5mm^2^), and kidney (0.5mm^2^). Entire atria imaged in place of fields of view (variable size). <u>Cardiac lymphatic localization:</u> Peri-lymphatic area was designated as an area within 25μm of LYVE1^+^ regions. 4-8 lymphatic vessels identified per myocardium. <u>Central/Portal Vein localization (CV, PV):</u> Cellular localization to the central or portal veins was designated as any cell within 50 μm of a CD31^+^ CV or PV. 2-8 regions designated as a CV; 5-16 regions designated as a PV acquired per animal. <u>Renal vessel association:</u> One large vessel was analyzed per animal. Vascular associated was designated as any cell within 50µm of a CD31^+^ region, measuring 0.03-0.24mm^2^

#### Macrophage sensitivity to cell death

<u>Bone Marrow-Derived Macrophage generation:</u> Bone marrow was harvested from femur and tibia from 16-week WT and STING^-/-^ mice. Bone marrow was cultured and differentiated for 7 days in RPMI with 30% L929 supernatant, 20% FBS, 2% Pen/Strep. <u>Isolation of unelicited peritoneal macrophages:</u> Resident peritoneal macrophages (unelicited) were collected from 16-week-old WT and STING^-/-^ mice through intraperitoneal injection of 10mL ice cold PBS with an 18G syringe through the lower left quadrant. Mice were shaken 16 times before macrophage-enriched PBS was aspirated from the peritoneal cavity.

#### In vitro induction of cell death

Induction of macrophage cell death from 16-week-old WT and STING^-/-^ was quantified as previously described. In brief, bone marrow-derived or peritoneal macrophages were plated in DMEM at 10^6^ cells/mL in a 384-well plate (25µL). Cells were incubated for 4h prior to stimulation with LPS (100ng/mL), FasL (100ng/mL), zVad (50µM), Nigericin (1µM), H202 (1mM), 3-MA (2mM), or Etoposide (150µM) with propidium iodide (PI, 10ng/mL). As a readout for cell death, cellular uptake of PI was measured with the Cytation3 microscope every 30 minutes for 16 hours. 5 technical replicates were included per biological replicate.

#### Multiplexed Cytokine/Chemokine with Eve Technologies

Two mouse multiplex cytokine array panels were included to test serum for protein levels of the following chemokines/cytokines from young, 12mo, 18mo, 24mo WT and STING^-/-^ mice (n=4-5): Eotaxin, G-CSF, GM-CSF, IFNγ, IL-1α, IL-1β, IL-2, IL-3, IL-4, IL-5, IL-6, IL-7, IL-9, IL-10, IL-12

(p40), IL-12 (p70), IL-13, IL-15, IL-17A, IP-10, KC, LIF, LIX, MCP-1, M-CSF, MIG, MIP-1α, MIP-1β,

MIP-2, RANTES, TNFα, VEGF, (LIX results also included but not validated), EPO, 6Ckine, Fractalkine,

IFNB1, IL-11, IL-16, IL-20, MDC, MCP-5, MIP-3a, MIP-3B, TARC, TIMP-1. Analysis was conducted

using GraphPad Prism 9 (GraphPad Prism).

#### Single cell analysis of tissue associated immune and stromal cells

##### 10x#Genomics Preparation

All steps were performed according to 10X Genomics standardized protocols following the generation of single cell suspensions. Young and aged samples from each organ and genotype were processed with 10x Chromium Single Cell 3’ Gene Expression platform (10x Genomics). Samples were loaded onto the 10X Chromium Controller to generate Gel Beads-in-emulsion (GEMs) and cDNA with cell-specific barcodes and libraries for single cell RNA-sequencing experiment one and two, v2 and v3 Chemistry, respectfully. Sample- and cell-specific libraries were sequences on Illumina NextSeq 550 (Tufts Medical School) targeting a depth of 75,000 reads per cell (experiment one and two).

##### Alignment / Preprocessing

Raw sequencing results from NextSeq 550 in BCL format was used as input for basecalling, demultiplexing and fastq file generation were performed using cellranger (v3.0.2) mkfastq pipeline. FASTQ files were aligned to the mm10 transcriptome, filtered, and quantified (barcode and UMI), using cellranger count, before all applicable datasets were aggregated using cellranger aggr using the recommended pipeline variables.

##### Preprocessing – Combined tissue dataset

All applicable samples from three tissues (heart, liver, kidney) were integrated using cellranger aggr, as above. The Seurat package (v4.1.1) ^82,83^ was used for pre-processing, normalization, dimensionality reduction, and cluster identification for all tissues as a singular dataset. The combined dataset was filtered for genes found in ≥ 3 cells and cells with ≥ 200 features and ≥ 500 total reads.

##### Preprocessing – Individual tissues

Pre-processing, normalization, dimensionality reduction, and cluster identification were performed with the Seurat package, as above. Dataset pre-processing and cell type identification were computed for each individual tissue. Similarly, individual datasets (2/24mo and 3/12mo batches) were separately pre- processed for initial clustering and cell typing. Batches were filtered for genes represented in ≥ 3 cells, and cells with ≥200 quantified features and ≥ 500 total read counts. Batches then underwent initial clustering at which point clusters were removed that were primarily characterized by high mitochondrial content as described below.

##### Clustering and cell type identification of scRNAseq data

Expression counts were normalized using the Seurat base function SCTransform(), including percent mitochondrial content and total read counts as variables to regress (vars.to.regress). Normalized values underwent principal component dimensionality reduction through the Seurat RunPCA() function, at which point the number of informative principal components was determined by their relative standard deviation, using the Seurat ElbowPlot() function. Each sample in individual batches was then integrated using the Harmony R package (v0.1.1) ^84^ using the RunHarmony() function. After dimensional reduction and integration, for the individual tissue datasets, clusters were identified using the Seurat functions, FindNeighbors() and FindClusters(). Clusters characterized primarily by high mitochondrial content in individual tissue datasets were identified using the Seurat function VlnPlot() and removed. This clustering procedure was repeated on the remaining dataset until cells were no longer primarily identified by high mitochondrial content. Uniform Manifold Approximation and Projection (UMAP) was used to visually represent cluster-based analyses, using the Seurat RunUMAP() function. Cell type assignments of clusters were characterized by differential expression analysis between cluster profiles, using the Seurat FindConservedMarkers() function for individual datasets, and FindMarkers() function for the combined dataset.

##### Integration of scRNAseq batches

To increase effective resolution of the scRNAseq datasets for determining cell type clusters from individual tissues, the two batches per tissue were combined and the cells were clustered with same workflow for initial clustering described above. To increase effective resolution of determining tissue macrophage populations, these steps were subsequently performed on macrophages from each tissue. For characterization of differences between cell types and samples and account for any batch effects, additional pre-processing of the integrated datasets were performed as follows. Genes previously reported to be primarily indicative of sample quality (mitochondrial genes, ribosomal genes, Gm42418, AY036118) ^85^ were removed and samples were re-normalized using SCTransform() and re-combined using the Seurat function PrepSCTFindMarkers(). Finally, samples were de-noised and low-transcript expression was recovered through the R package Rmagic (v0.1.0) ^86^ function, magic().

##### Differential Gene Expression Analysis

Tissue fingerprinting: Marker genes for each sample were identified in an age and tissue dependent manner and filtered for adjusted p-values < 0.1 and log2 fold change > 0.2. The average expression of each gene per cluster was then calculated per sample using the Seurat function AverageExpression(). Average Expression: Differentially expressed genes (DEGs) were identified in an age-dependent manner for each identified cluster and filtered for adjusted p-values < 0.1 and log2 fold change > 0.2. The average expression of each gene per cluster was then calculated per sample (age/genotype) using the Seurat function AverageExpression().

Single-cell expression: Age-dependent and upregulated DEGs were calculated per each representative cell type, filtered for significance (adjusted p-value < 0.1), and ranked based on logs fold change. Top DEGs

1. (25) were then filtered and used to generate single cell heatmaps using the Seurat function DoHeatmap() and relative counts were extracted using the Seurat function FetchData().

##### Pathway analysis

DEGs were calculated using the Seurat standard FindMarkers() function. In brief, all genes positively enriched in experimental clusters were calculated with a log2 fold change threshold of 0.2. DEGs were then fed into an automated pathway analysis pipeline (grprofiler2) to calculate enriched gene ontology (GO) pathways. GO pathways were filtered for “GO:Biological Processes” with a term size < 2500 genes and more than 3 genes contained in the DEG list. As gene were previously designated as significantly differentially expressed between groups, no further significance thresholding was performed. Enriched pathways were sorted by term “recall” (proportion of genes in the dataset to the size of the GO term). For broad investigation of enriched pathways in Figure 2A: all enriched GO terms were fed into a clustering algorithm (Revigo) ^87^ to generate “representative terms” based on “semantic similarity” between GO pathways.

##### Immune signaling induced by serum immunokines

Characterization of relative (theoretical) cytokine activity was performed with the R package NicheNet (v1.0.0). ^13^ In brief, NicheNet file human gene symbols were converted to murine symbols and Rmagic processed gene expression values were quantile scaled and utilized to estimate relative ligand activity using the NicheNet functions convert_human_to_mouse_symbols(), scale_quantile(), and predict_single_cell_ligand_activities(). Mean Pearson correlation scores were calculated for cytokine activity for each cell type in the 24mo WT sample. Top results were identified by: a mean activity score > 0.05 in at least one cell type, and cell types with a mean activity score > 0.05 in at least one ligand.

##### Single Cell Gene Set Variation Analysis (GSVA)

To characterize differential pathway activity between cells and samples, a gene set projection of gene expression profiles was performed with the R package Gene Set Variation Analysis (GSVA, v1.42.0) ^28^. In brief, Murine gene sets annotated to biological processes were obtained from Gene Ontology (GO) ^88^ and Kyoto Encyclopedia of Genes and Genomes (KEGG) ^89,90^ with the R package gage (v2.44.0)^91^ using the functions go.gsets() and kegg.gsets(). Murine entrez IDs were mapped to murine gene symbols with the R package pathview (v1.34.0)^92^ using the function geneannot.map(). In addition, we included curated pathways for inflammation, interferon stimulated genes^17^ and senescence pathways (GO, KEGG).

Prior to running GSVA, genes that were not quantified in at least one sample for the given cell type were removed, and murine gene sets with fewer than 25% of their genes and fewer than 25 total genes in these filtered data were removed. GSVA activity scoring was performed on log2-transformed Rmagic processed gene expression values independently for each batch and cell type. For each cell type, differential analysis of GSVA activity scores between samples was performed with linear modeling using the R package LIMMA (v3.50.1) ^93^. Genotype-stratified statistical testing of changes in mean GSVA activity scores between ages, as well as differences in these changes between genotypes, was performed using the LIMMA function makeContrasts(). Multiple hypothesis correction of nominal p-values was performed using Benjamini-Hochbern false discovery rate (FDR)^94^ and statistical significance was established based on contrasts yielding an FDR adjusted p-value < 0.05.

##### Relative spatial interactions

The RNAMagnet package ^23^ was used to predict physical interactions (RNAMagnetAnchors) between selected cells based on relative expression of receptor:ligand pairs. All appropriate stromal cells were included as potential “anchors” and interaction scores were calculated for all clusters of the dataset.

Interaction scores were ranked to determine the most likely interaction between cell types. Primary interactions were visualized as relative proportion of cell clusters to understand age- and genotype-associated changes of predicted cellular localization.

##### Immune signature enrichment and dysregulation

GSVA activity scores were linear transformed to a positive integer scale. Using the young WT cell clusters as a control, all single cell activity scores for INF, ISG, Sen were normalized to the mean pseudocount-transformed GSVA activity score of the respective cell cluster to generate relative pathway enrichment for each sample and cell cluster. All cells with normalized GSVA scoring above 1.2x the mean of the young WT cell cluster (heart, kidney; 1.1x for the liver) were considered as “positively enriched” for each specific pathway. Cells with normalized GSVA activity scores greater than the indicated threshold for two or three independent pathways were considered double or triple positive cells.

##### Quantification and Statistics: Single Cell RNA sequencing

Differentially expressed genes were calculated with Seurat built in functionality (FindMarkers, FindAllMarkers) and p values were calculated with a Wilcoxon Rank Sum test. Significant genes were thresholded with a Bonferroni adjusted p value < 0.1.

### Survival Analysis

Healthy WT and STING^-/-^ mice of varying ages were designated to be included in survival analysis. Mice were monitored daily by the Division of Laboratory Animal Management (DLAM) for outward signs of disease, including daily scoring of body condition^95^. Body condition was only noted if mice progressed

+/- 1 relative unit. Date was recorded upon natural death or requirement of euthanasia per DLAM instruction for animal welfare. Birth, death date, body weight, and body condition score (if applicable) were recorded. Individual lifespan calculations were graphed in GraphPad Prism, statistical testing was performed using the Mantel-Cox Logrank test. Statistical significance of survival analysis testing was considered at p-value < 0.05.

#### Quantification and Statistics

All data are shown as mean ± SEM. Sample (n) indicates total number of samples included in the analyses and multiple experiments. A Wilcoxon rank sum test was used for all single cell differential gene expression analyses. Data thresholding was limited to Bonferroni corrected p value < 0.1 as indicated.

One-way ANOVA with Tukey correction was performed where indicated to test either age (WT only) or genotype (WT and STING^-/-^) effects on sample distribution. A Two-way ANOVA was performed for TUNEL assays to test combined age and genotype effects. Power analyses were not performed prior to experimental design. GraphPad Prism (v9) and built-in Seurat functions were used to calculate statistical significance. All graphing and data visualization was performed in GraphPad Prism or with R. Statistical significance was defined as p < 0.1 (gene expression analysis, adjusted) or p < 0.05 (all other analyses, adjusted).

**Supplemental Information 1: Curated gene lists for primary immune terms**

**Supplemental Information 2: DEGs mapping to unique gene ontology pathways as related to Figure 2A**

---

**Supplemental Figure 1:**
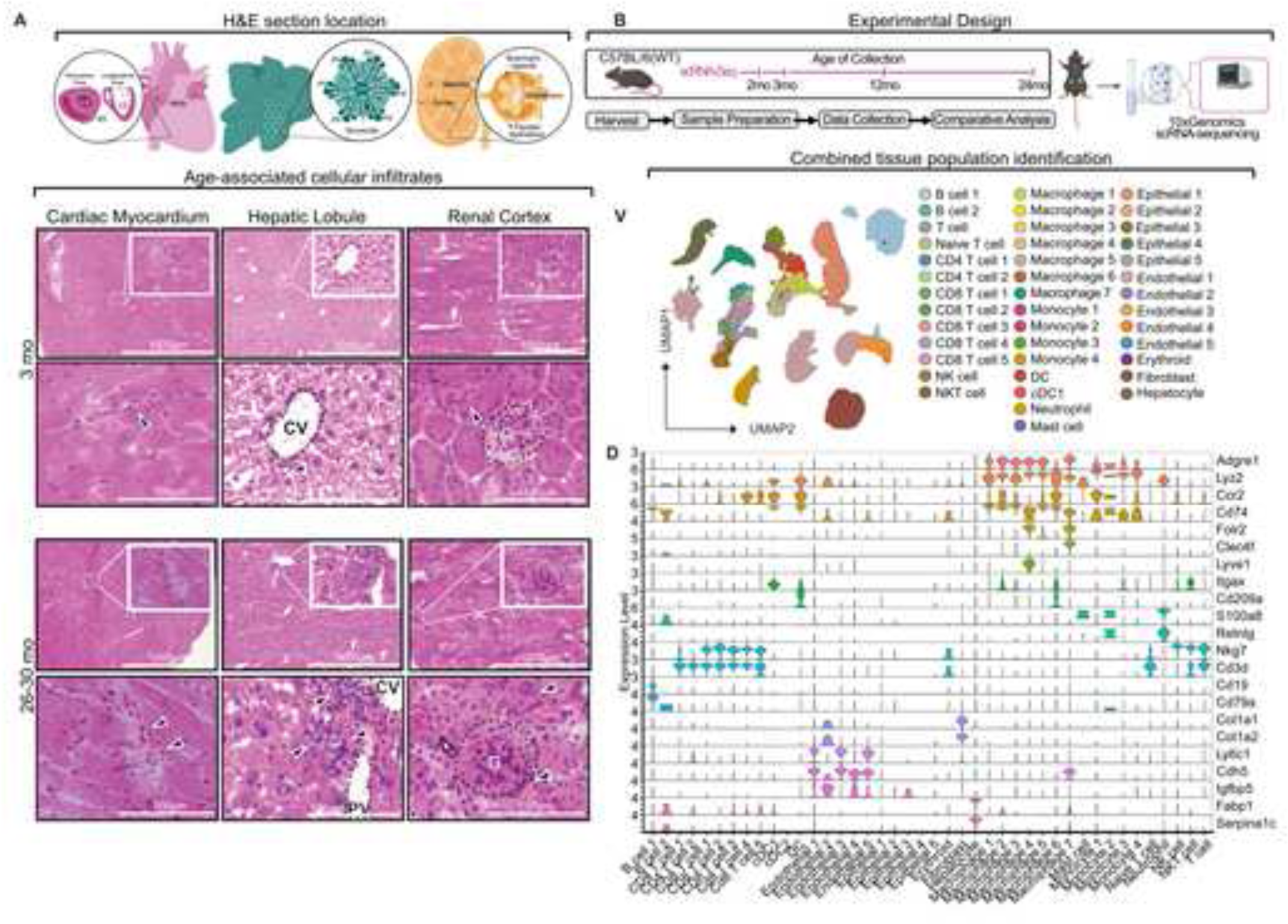
Aging drives tissue inflammation distinct from circulating inflammatory effectors. (A) Section location for H&E histopathology in young (3mo) and aged (26-30mo) WT heart (left, cardiac myocardium), liver (middle, hepatic lobule), and kidney (right, renal cortex). Black arrows indicate regions of cellular infiltration. CV; central vein, G; glomerulus. **(B)** Experimental design for single cell RNA sequencing (scSeq) preparation and analyses. **(C)** UMAP of heart, liver, and kidney scSeq datasets highlighting samples by age, lineage, and transcriptionally distinct clusters of cells. Cell typing as annotated. **(D)** Violin plots quantifying relative expression of common lineage identifying genes used in part for cell typing.

**Supplemental Figure 2:**
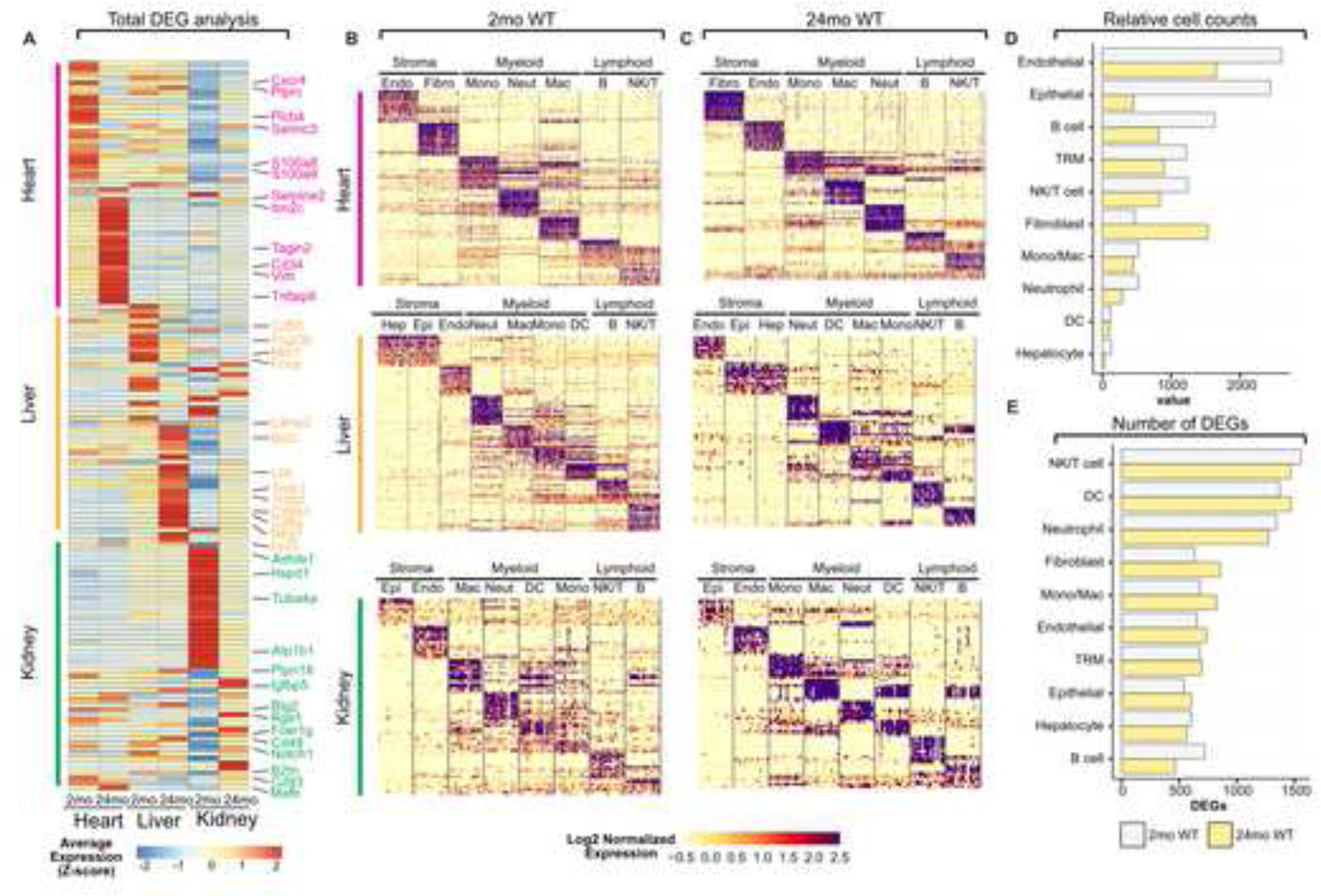
Aging tissues are imprinted with unique transcriptional signatures. **(A)** Relative average expression of top marker genes from 2mo and 24mo WT tissues (Magenta; heart, gold; liver, green; kidney). **(B-C)** Single cell heatmap of top marker genes per cell grouping in 2mo **(B)** and 24mo **(C)** WT heart (magenta), liver (gold), and kidney (green). Highly expressed genes in purple, lowly expressed genes in light yellow (Fibro, fibroblast; Endo, Endothelial; Epi, epithelial; Hep, hepatocyte; Mono, monocyte; Mac, macrophage; Neut, neutrophil; DC, dendritic cell; B, B cell; NK/T, NKT cell/T cell). **(D)** scRNAseq cell counts from 2mo (grey) and 24mo (yellow) samples independent of tissue. **(E)** Total number of DEGs calculated as marker genes per cell group in 2mo (grey) or 24mo (yellow) combined tissue samples. Statistically significant genes determined with a Wilcoxon rank sum test with Bonferroni adjusted p-value < 0.1.

**Supplemental Figure 3:**
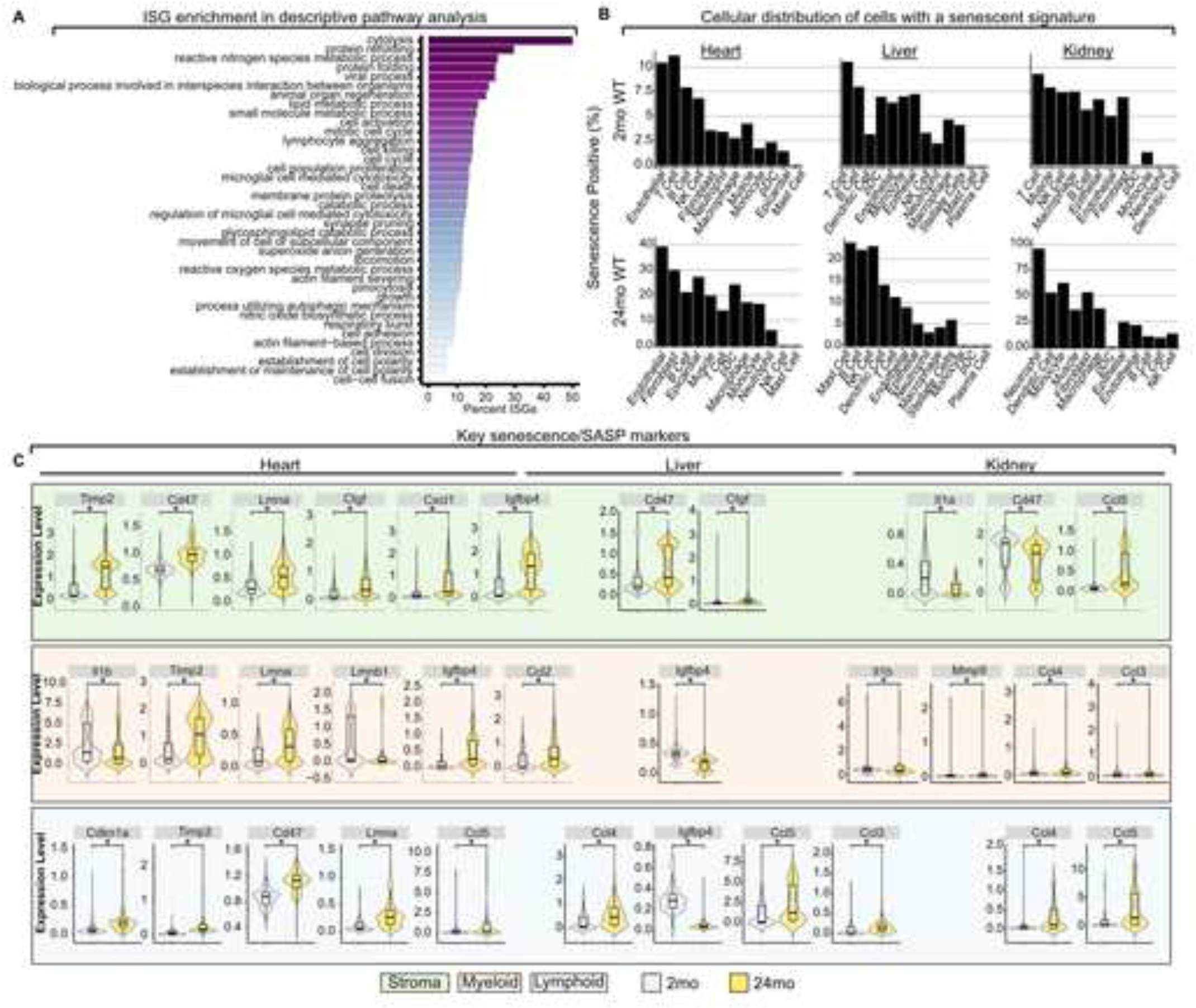
Senescence signatures are not temporally regulated in aging tissues. **(A)** Enrichment of ISGs as a proportion of total genes within summary GO pathways enriched in aged cell types from Figure 2A. **(B)** Bar graphs representing percent of cells relatively enriched in a senescence signature as calculated by GSVA in 2mo and 24mo WT heart (left) liver (middle) and kidney (right). **(C)** Key senescence associated and SASP genes significantly altered with age (Bonferroni adjusted p value < 0.05) in stromal (green box), myeloid (red box) and lymphoid (blue box) cells from heart (left), liver (middle), and kidney (right). Statistically significant genes determined with a Wilcoxon rank sum test. \**p ≤ 0.05*.

**Supplemental Figure 4:**
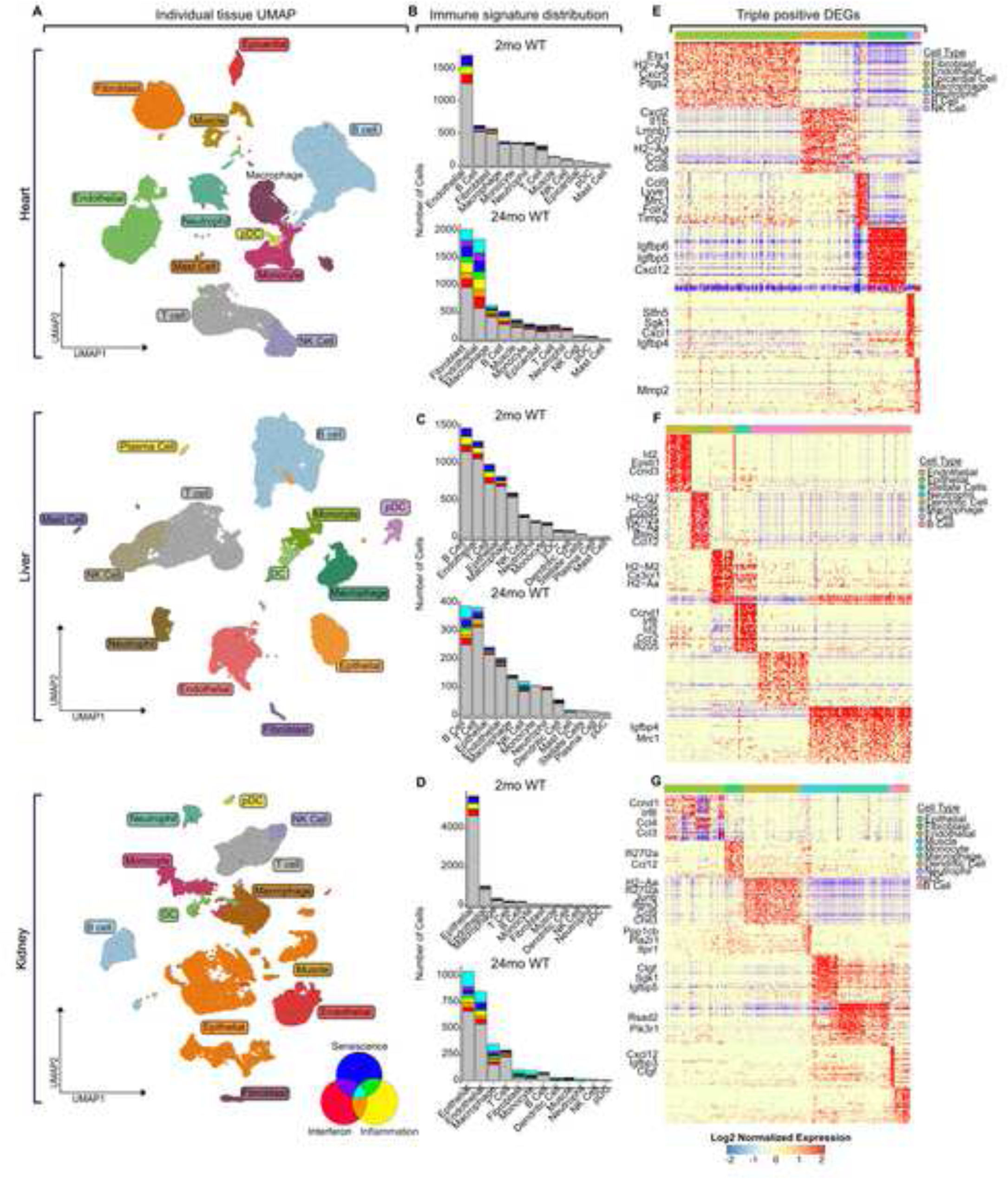
Aged triple positive cells maintain tissue imprinting. (A) UMAP projections of single tissue datasets highlighting unique cell groupings. **(B-D)** Stacked bar graph of 2mo (top) and 24mo (bottom) heart **(B)**, liver **(C)**, and kidney **(D)**. Bar graphs represent raw counts of cells that are enriched in immune signatures, including double and triple positive cells from unique cell groups. **(E-G)** Single cell gene expression profiles of triple positive immune cells from 24mo heart **(E)**, liver **(F)**, and kidney **(G)**. Statistically significant genes determined with a Wilcoxon rank sum test with Bonferroni adjusted p-value < 0.1.

**Supplemental Figure 5:**
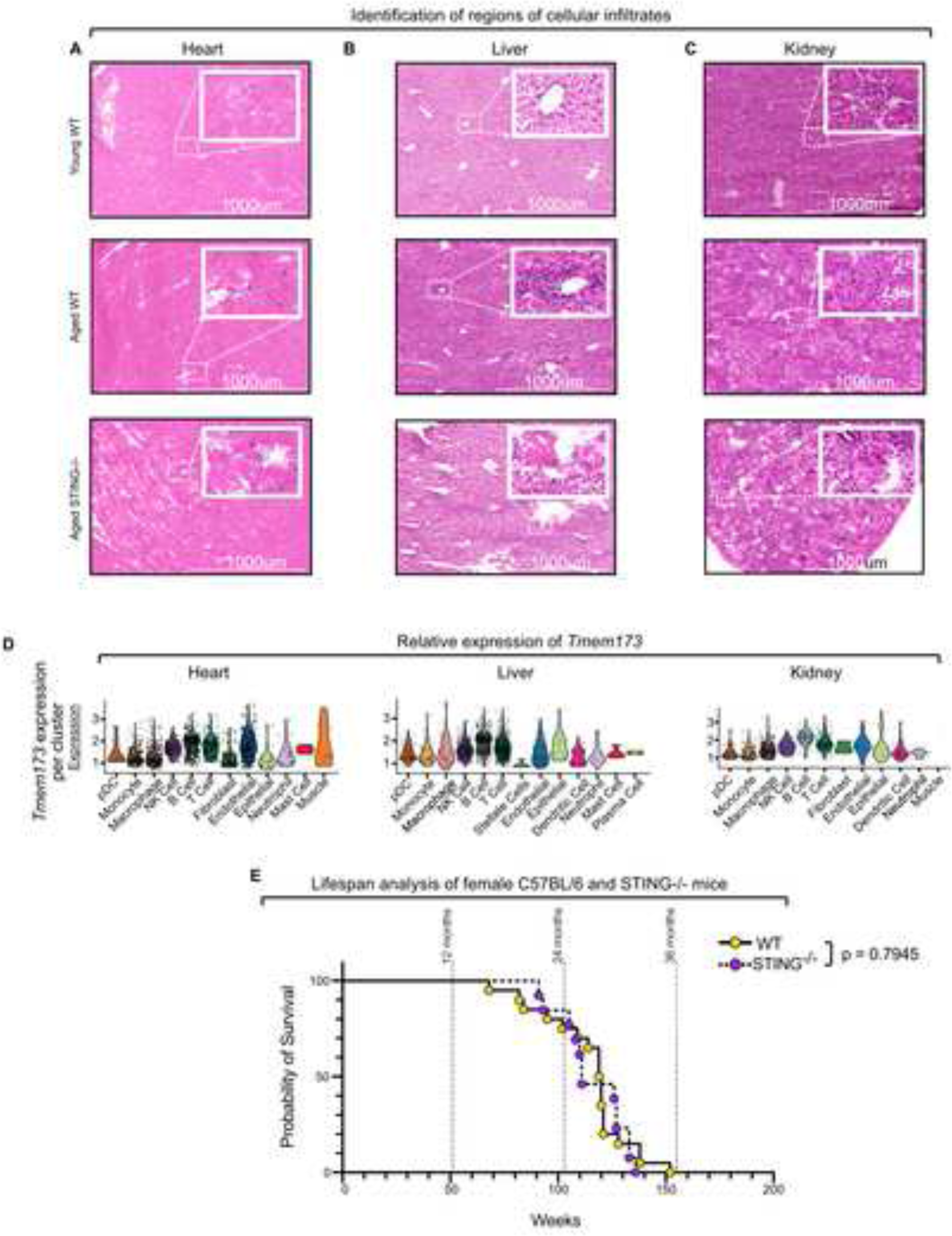
STING is sufficient to restrain cellular inflammation and promote homeostasis in male mice. (A-C) Macro view of H& staining in Figure 3B-D in young (3mo) and aged (26-30mo) WT and STING^-/-^ heart **(A)**, liver **(B)**, kidney **(C)**. **(D)** Relative expression of STING (*Tmem173*) in stromal, myeloid, and lymphoid cells from the heart (right), liver (middle) and kidney (left). **(E)** Kaplan-Meier survival probability curves comparing WT (solid line, yellow circles) and STING^-/-^ (dashed line, purple circles) females. Statistics were calculated with a Mantel-Cox Logrank test.

**Supplemental Figure 6:**
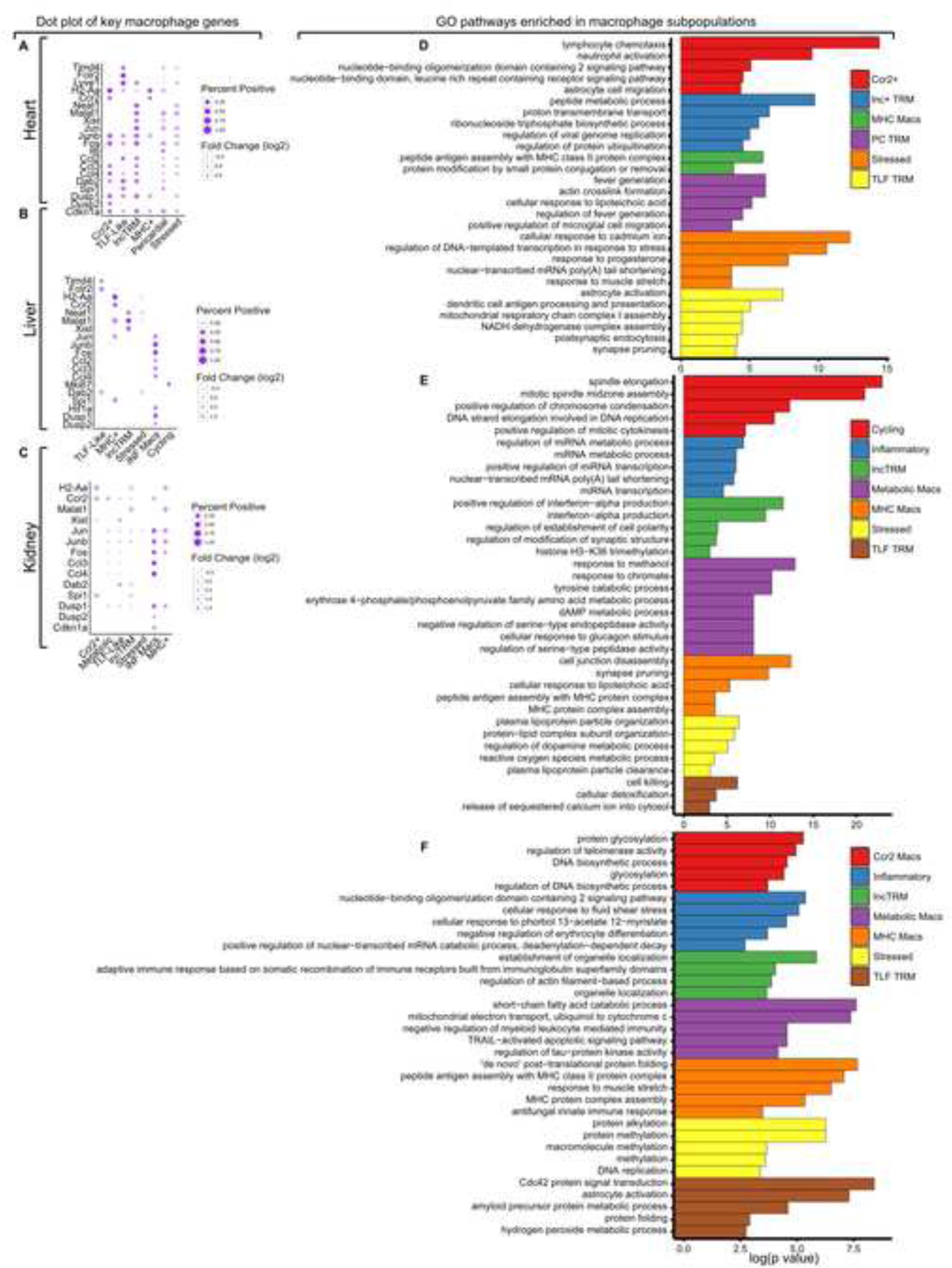
Identification of resident macrophage subsets. (A-C) Dot plot of top genes from transcriptionally distinct macrophage subsets for cluster identification in the heart **(A)**, liver **(B)**, and kidney **(C)**. **(D-F)** Gene ontology pathway analysis of DEGs from each transcriptionally distinct macrophage subset in the heart **(D)**, liver **(E)**, and kidney **(F)**. Macrophage populations annotated to the side. GO pathways scaled by log(p value). Statistically significant genes determined with a Wilcoxon rank sum test with Bonferroni correction were provided for GO pathway enrichment.

**Supplemental Figure 7:**
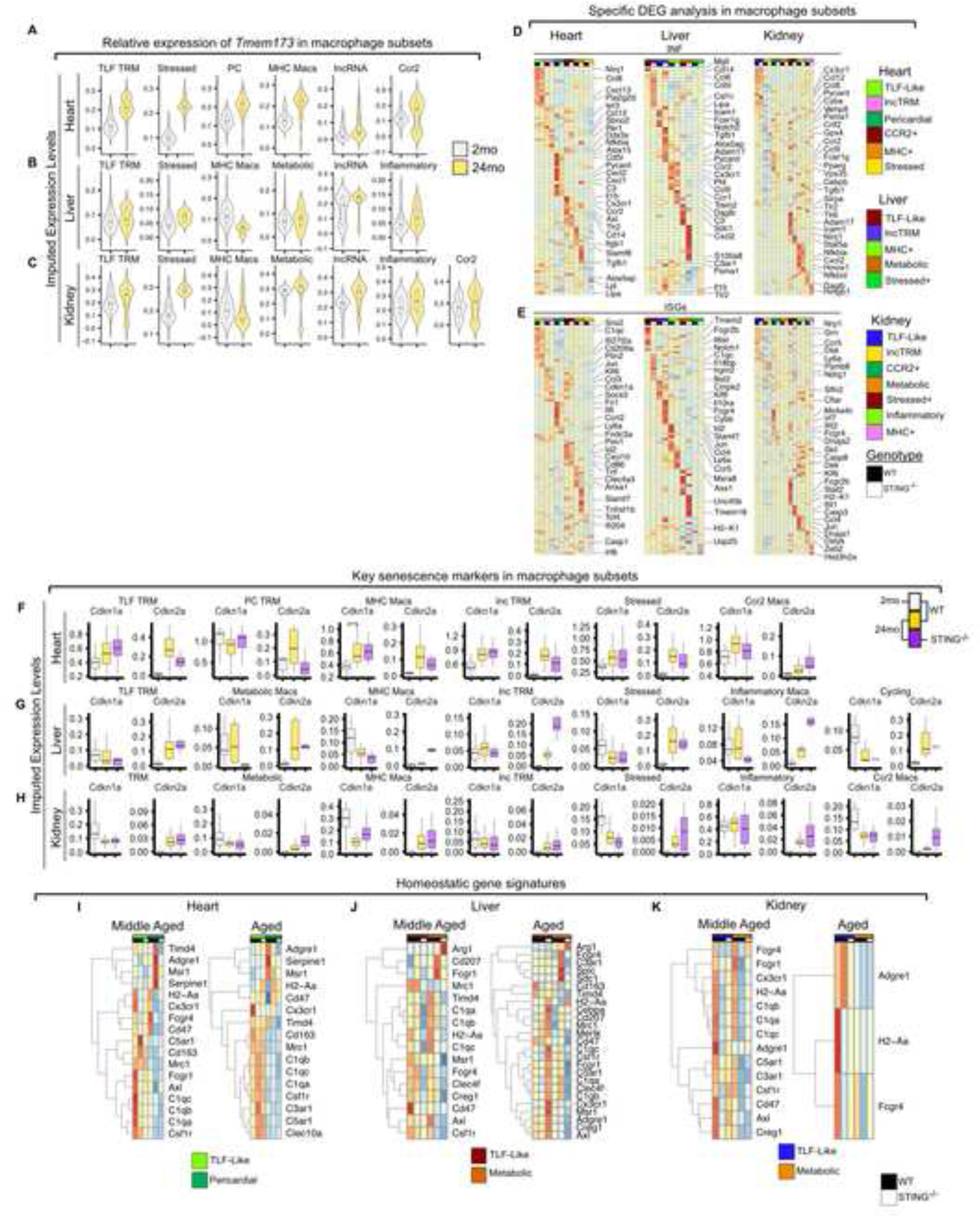
STING regulates initiation of senescence and homeostatic capability of tissue macrophages. (A-C) Violin-box and whisker plots of STING expression in tissue macrophage subsets in 2mo WT (grey) and 24mo WT (yellow) from the heart **(A)**, liver **(B)**, and kidney **(C)**. **(D-E)** Top inflammatory **(D)** and ISG **(E)** genes from the heart (left), liver (middle), and kidney (right). Genotype and macrophage subtype as annotated above. **(F-H)** Relative expression of key senescence marks from macrophage subsets in the heart **(F)**, liver **(G)**, and kidney **(H)**. 2mo WT, grey; 24mo WT, yellow; 24mo STING^-/-^, purple. **(I-K)** Average expression of curated homeostatic gene signatures from middle aged (left) and aged (right) heart **(I)**, liver **(J)**, and kidney **(K)** resident macrophages. Statistically significant genes determined with a Wilcoxon Rank Sum test with Bonferroni correction with an adjusted p value limit of 0.1. \**p ≤ 0.05*

**Supplemental Figure 8:**
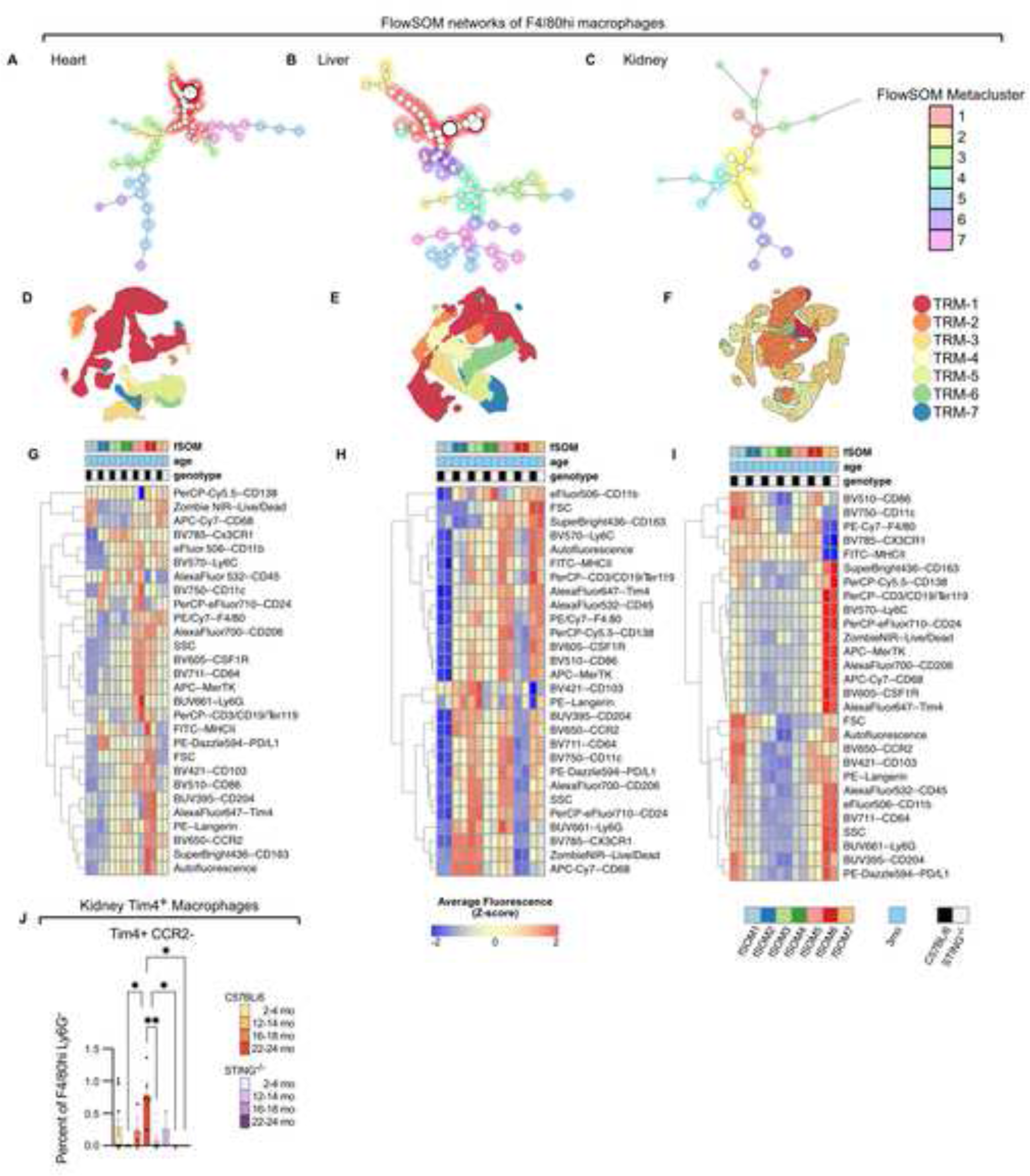
Tissue macrophages are comprised of distinct subtypes. (A-C) FlowSOM networks of F4/80^hi^ macrophages from heart **(A)**, liver **(B)**, and kidney **(C)**. Individual metaclusters color coded and annotated to the side. **(D-F)** UMAP of F4/80^hi^ macrophage with overlays of FlowSOM metaclusters from heart **(D)**, liver **(E)**, and kidney **(F)**. Individual macrophage populations color coded and annotated to the side. **(G-I)** Relative average expression of surface protein markers from young (2-4mo) F480^hi^ macrophages as measured by flow cytometry from heart **(G)**, liver **(H)**, and kidney **(I)**. Metaclusters, age, and genotype annotated above heatmap. **(J)** Proportion of Tim4^+^ F4/80^hi^ macrophages as percent of total F4/80^hi^ Ly6G^-^ cells from the kidney.

## Notes

### Competing Interest Statement

The authors have declared no competing interest.

