## Supplementary figures and images for "STING promotes homeostatic maintenance of tissues and confers longevity with aging"

### Supplemental Data 1

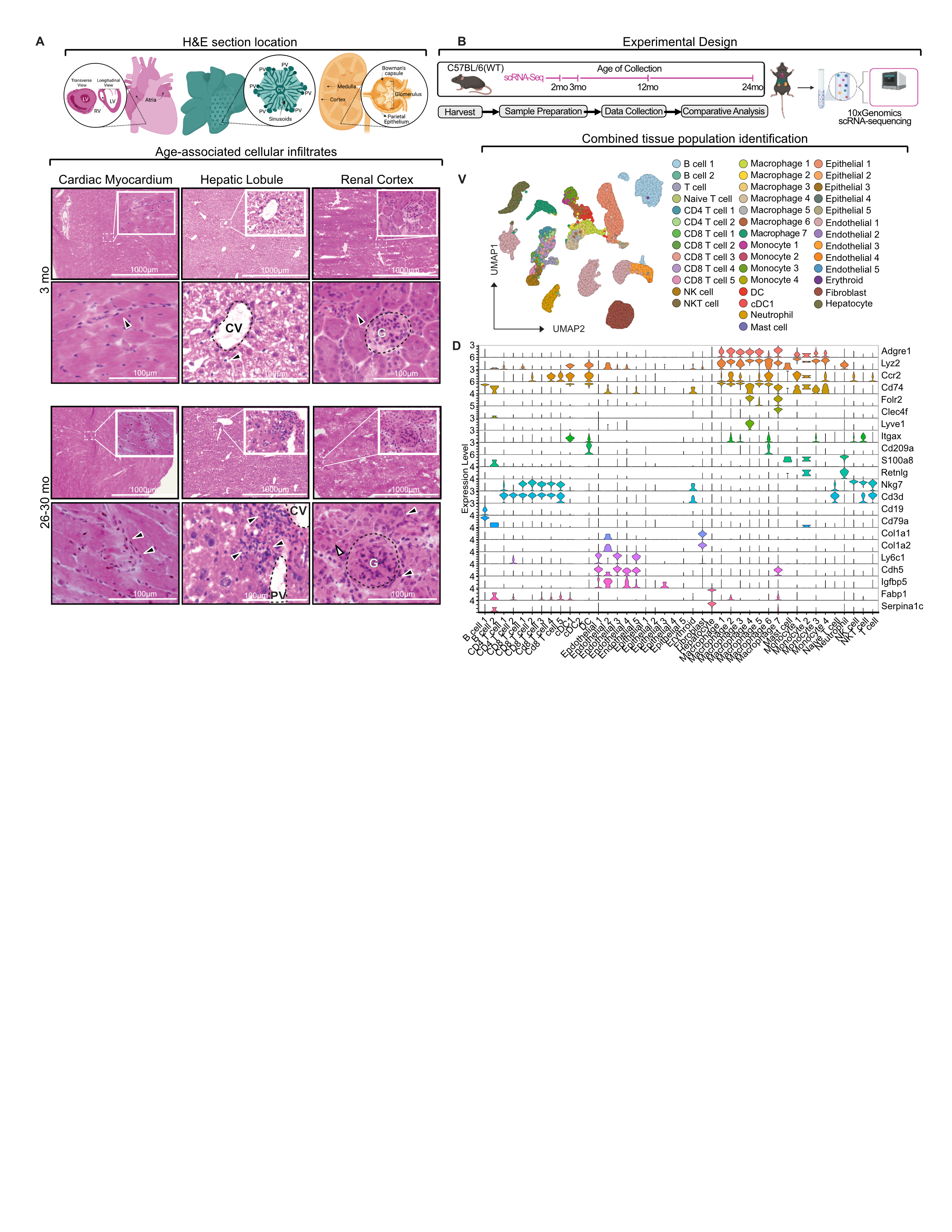

### Supplemental Data 2

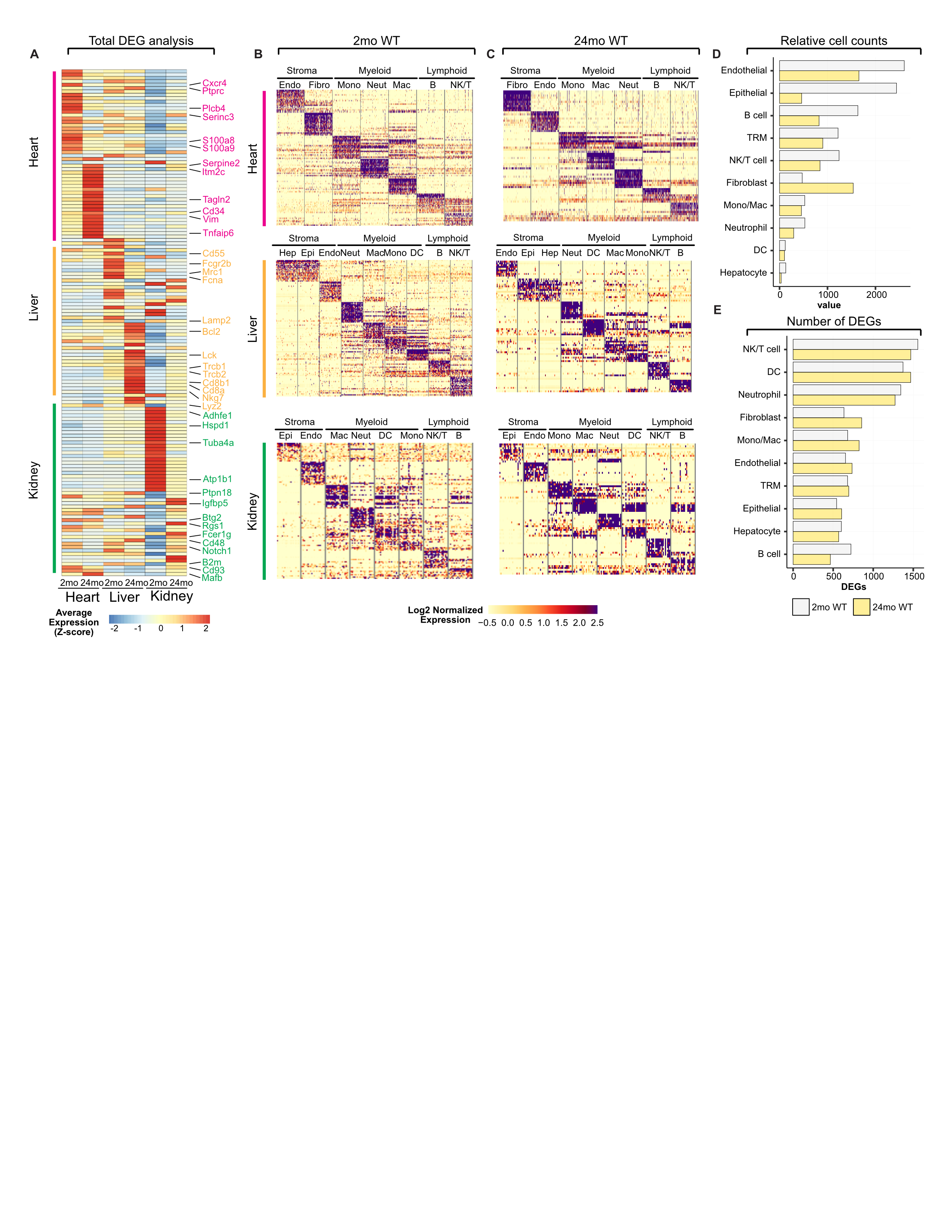

### Supplemental Data 3

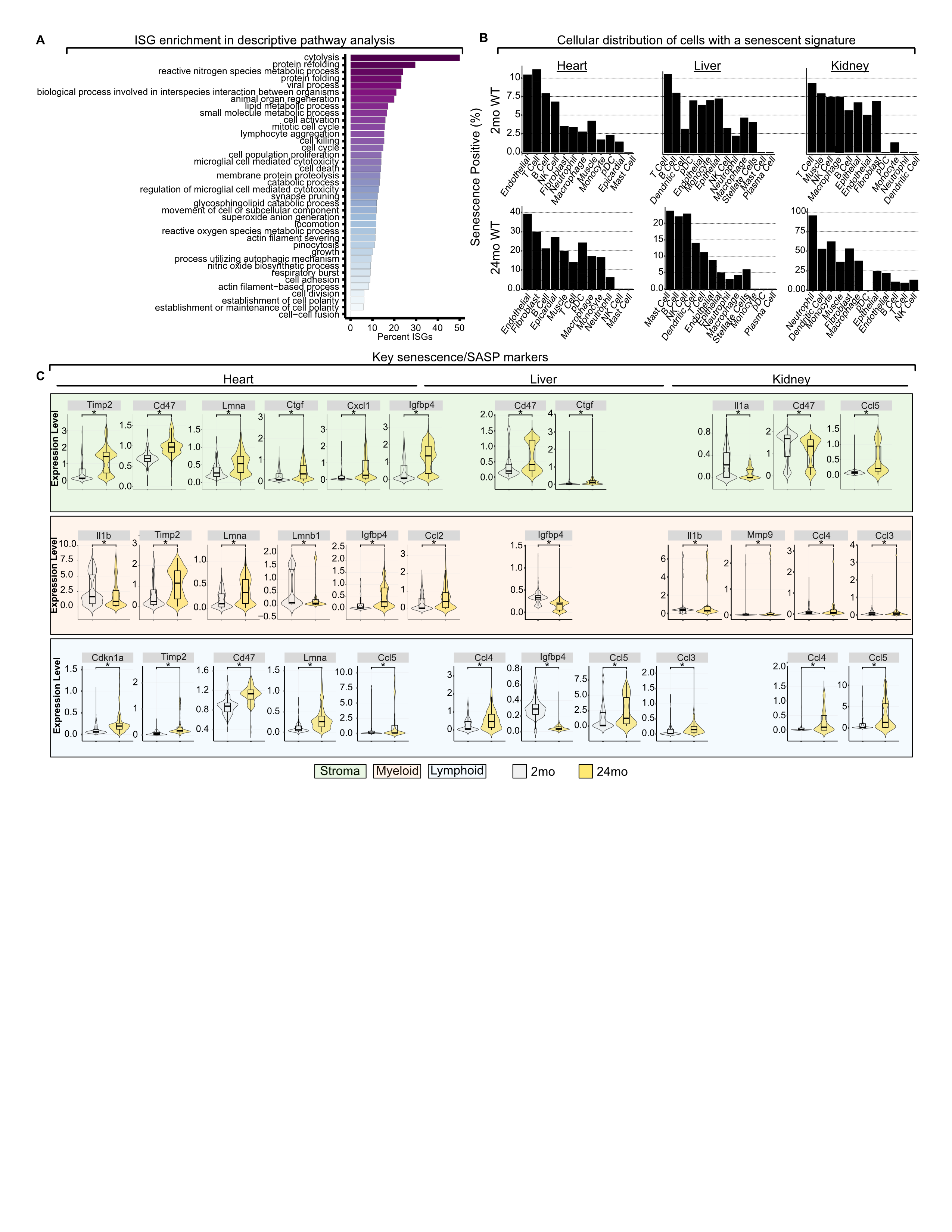

### Supplemental Data 4

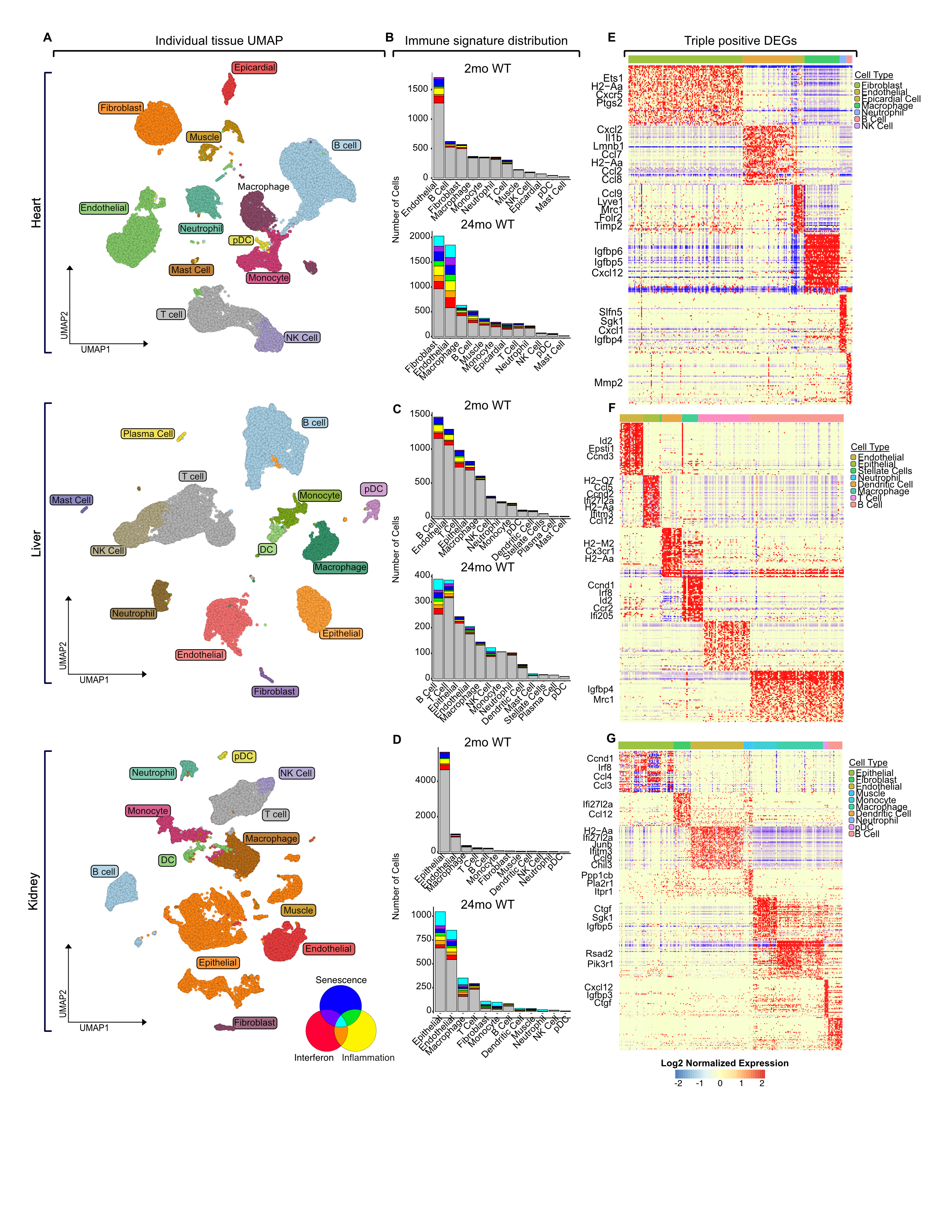

### Supplemental Data 5

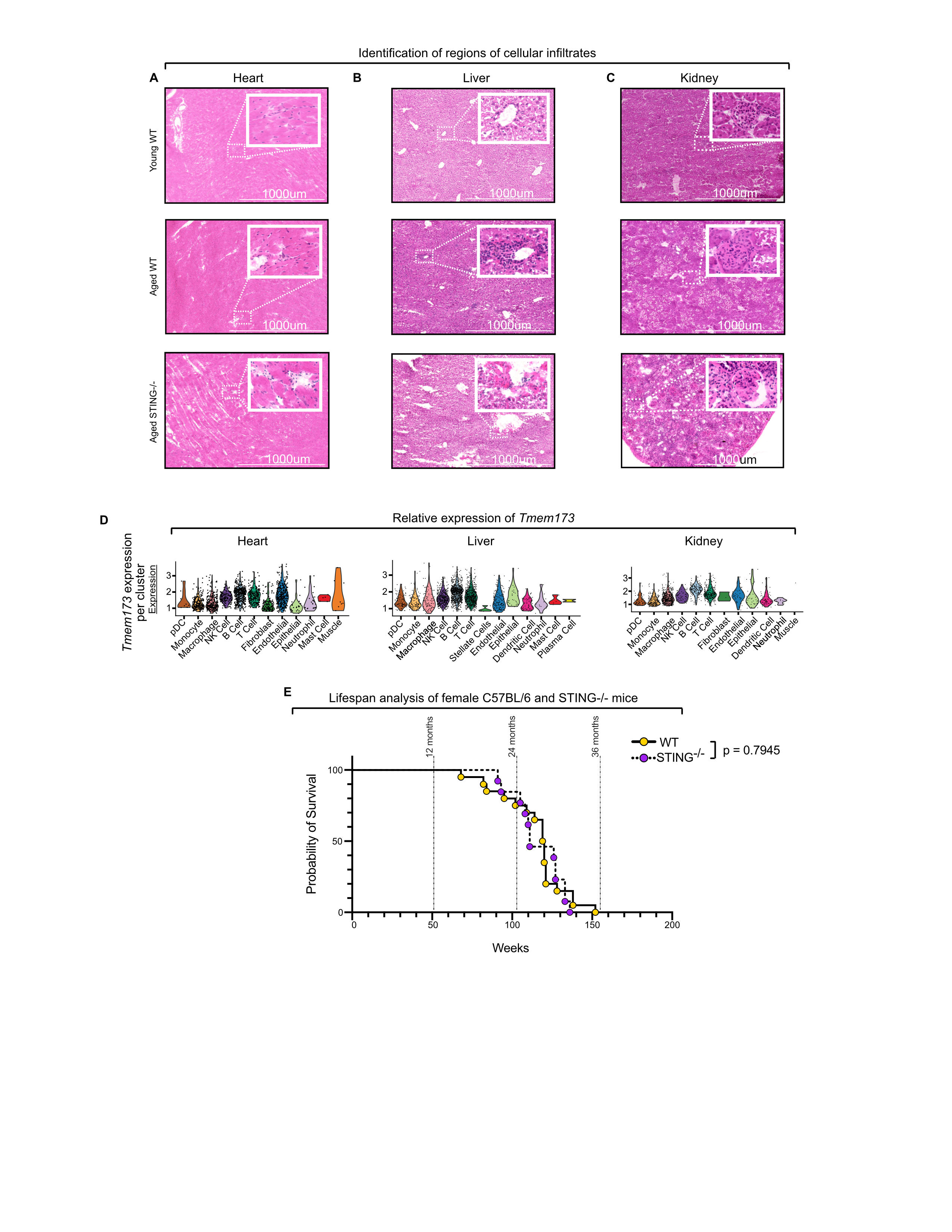

### Supplemental Data 6

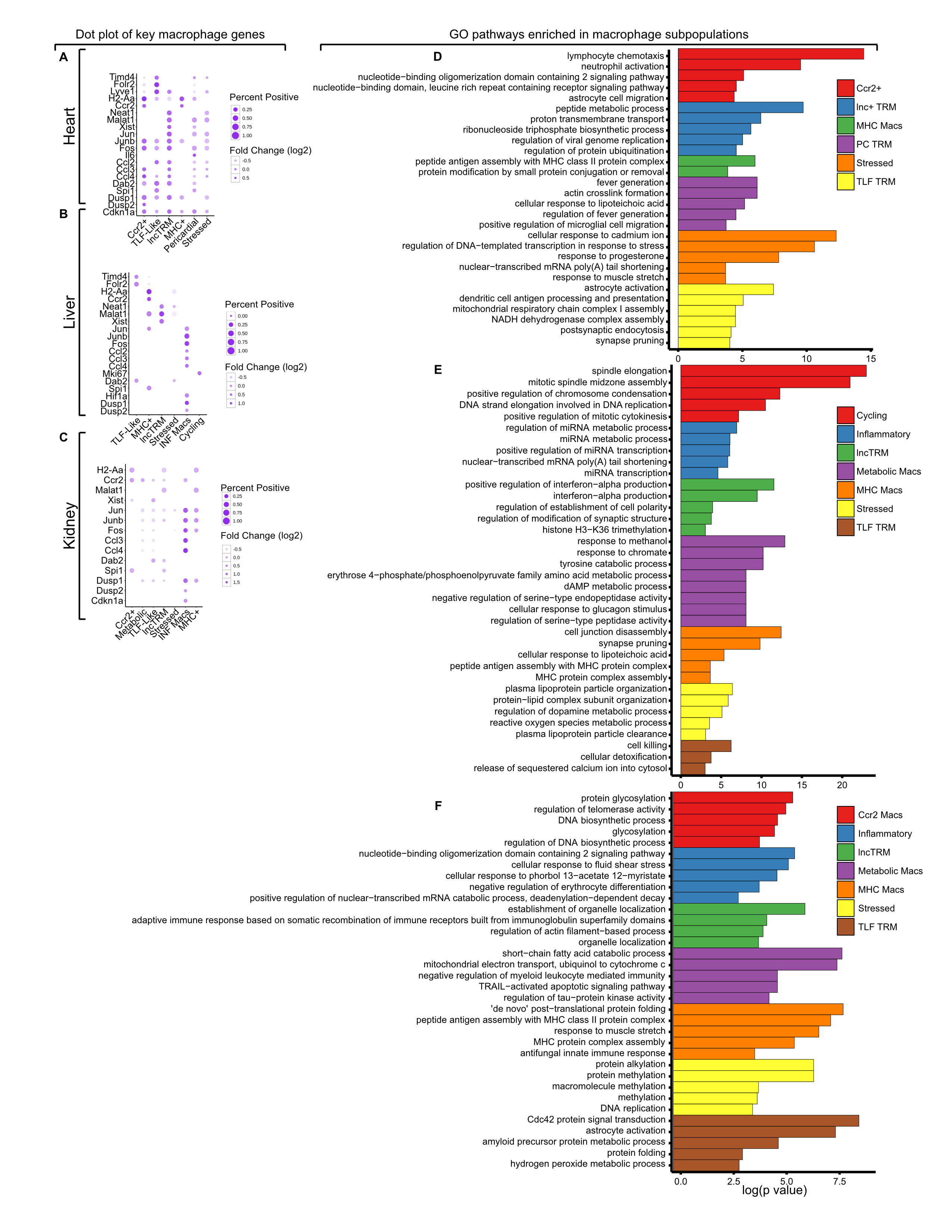

### Supplemental Data 7

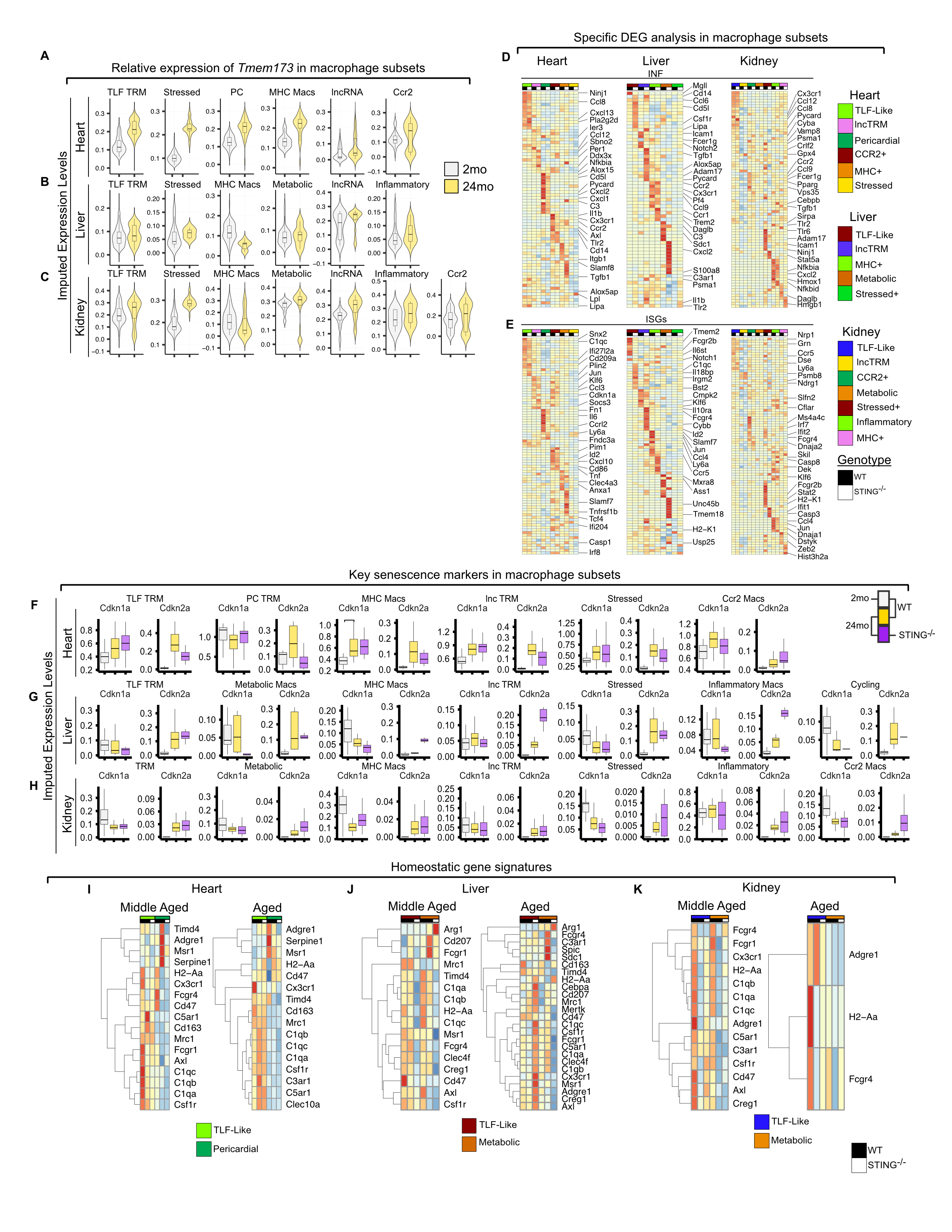

### Supplemental Data 8

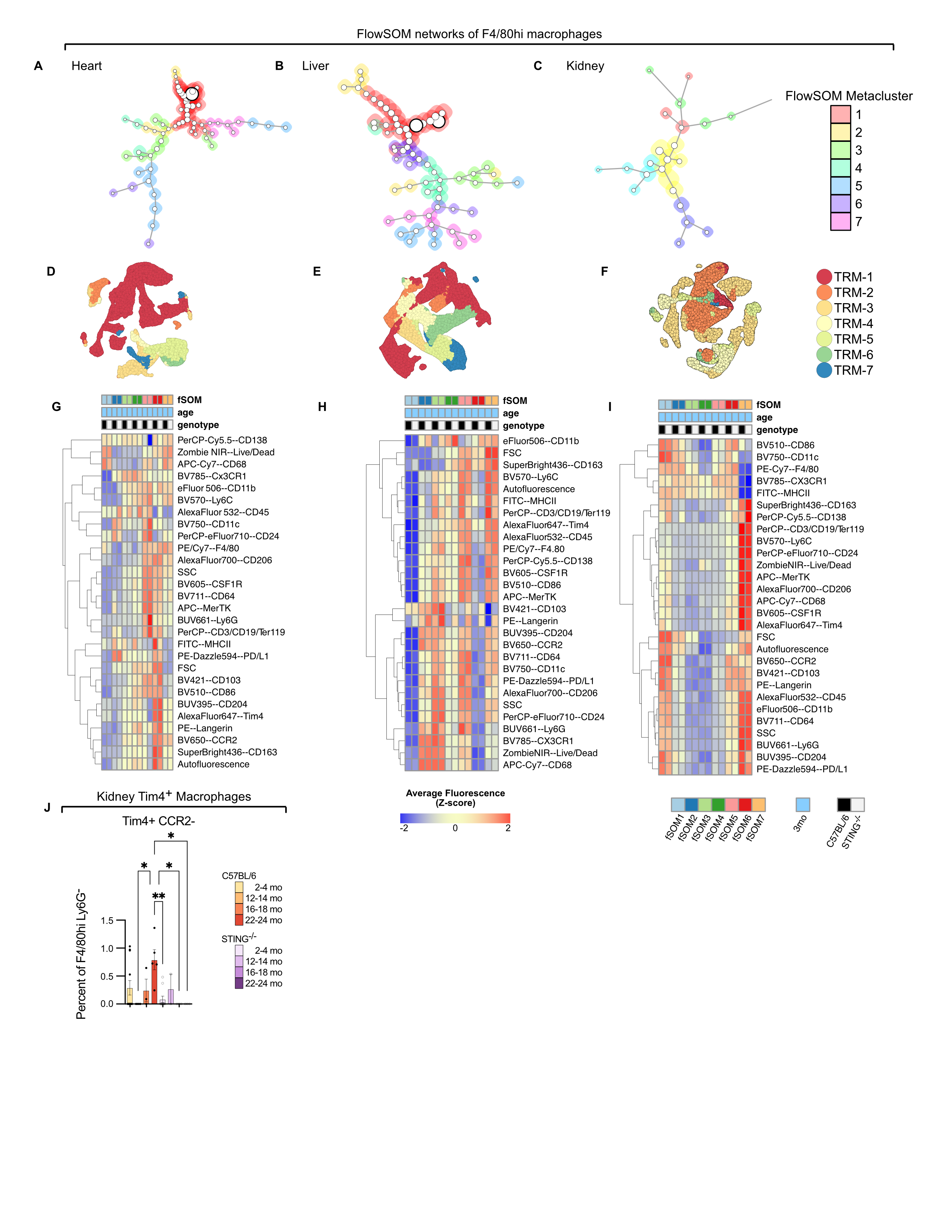
